# Cultivar-specific fungicide resistance emerges during a growing season in field populations of *Zymoseptoria tritici*

**DOI:** 10.1101/2024.08.07.606886

**Authors:** Firas Talas, Jessica Stapley, Bruce A. McDonald

## Abstract

*Zymoseptoria tritici* causes the most damaging wheat disease in Europe, septoria tritici blotch (STB). In Europe, STB is controlled mainly by fungicides and fungicide resistance is frequently reported. While fungicide resistance is thought to emerge mainly from standing genetic variation within field populations of *Z. tritici*, few studies have attempted to quantify the degree of fungicide resistance occurring at the field scale and to measure changes in frequencies of resistant strains following fungicide applications during a single growing season. Even fewer studies have considered the effects of different wheat cultivars on the emergence of fungicide resistance. We measured EC50 values for 1005 strains of *Z. tritici* sampled at two time points from 17 different wheat cultivars growing in a replicated field experiment that was treated with combinations of five different fungicides three times during the growing season. We found that field populations of *Z. tritici* can maintain a very high diversity in fungicide sensitivity phenotypes despite three fungicide treatments, with as much diversity found within a single field during a single growing season as has been described across all of Europe over several years. Multidrug resistance to two or more fungicides was found in 18.9% of the tested strains. We discovered that wheat cultivars that were more resistant to STB tended to be colonized by *Z. tritici* strains that exhibited higher fungicide resistance. We also found that specific wheat cultivars selected for resistance to specific active ingredients. Overall, our findings illustrate the many challenges associated with designing fungicide treatment programs that aim to reduce selection for fungicide resistance when confronted with a pathogen like *Z. tritici* that has a very high evolutionary potential.

## Introduction

*Zymoseptoria tritici*, the pathogen causing septoria tritici blotch (STB), poses a substantial threat as a destructive wheat pathogen, especially in Europe where it typically causes yield losses of 5-10% despite the intensive use of fungicides and deployment of resistant cultivars^13^. Approximately 70% of all fungicide applications on wheat in Europe are aimed towards controlling STB ^50^. Not surprisingly, *Z. tritici* populations have evolved resistance to fungicides in all regions where they are regularly applied. The rapid emergence of fungicide resistance is facilitated by the high evolutionary potential of *Z. tritici*, which reflects its frequent sexual recombination, a high effective population size, and substantial gene flow across regional scales ^34^. A recent analysis ^35^ indicated that each hectare of a wheat field with a moderate degree of STB infection (∼40% of leaves infected) contains ∼8 million different pathogen genotypes that produce ∼5 trillion pycnidiospores during a typical growing season, with ∼70 million of these spores expected to encode mutations for fungicide resistance. These findings indicate that most fungicide resistance is likely to emerge from standing genetic variation in fields affected by STB.

A diverse array of fungicides with various modes of action are applied to wheat fields to control STB and other diseases. Sterol biosynthesis inhibitors (SBIs) target the synthesis of ergosterol, a vital component of fungal cell membranes. SBIs are classified into several subgroups based on their specific interaction sites across the ergosterol biosynthetic pathway. Demethylation inhibitors (DMIs) are the largest subgroup, targeting a C14-demethylase (erg11/cyp51). DMIs consist of several chemical subgroups, including triazoles (e.g., metconazole, epoxiconazole, and propiconazole) and triazolinthiones (e.g., prothioconazole). Another subgroup within the SBIs is the amines, which target a Δ^14^-reductase and a Δ^8^®Δ^7^-isomerase within the sterol biosynthesis pathway. Although this subgroup includes various chemical groups, including morpholines, piperidines, and spiroketalamines (e.g., spiroxamine), the term “morpholines” is commonly used to refer to the entire subgroup. Another important group of fungicides is the succinate dehydrogenase inhibitors (SDHIs), which target complex II of the mitochondrial respiration chain. SDHIs comprise various chemical groups, including pyrazole-carboxamides (e.g., bixafen) ^14^.

Several approaches can be used to evaluate fungicide resistance in vitro. A classical method involves mixing a single concentration or serial dilutions of a fungicide with growth media before pouring Petri plates, enabling calculation of the minimum inhibitory concentration (MIC) (eg., ^24^) or differences in growth rates among isolates (e.g. ^29^). Another technique is the disk diffusion method, where the inhibition diameter is measured in millimeters to compare different concentrations (adsorbed onto filter discs) or isolates ^31^. Measuring growth rates based on changes in optical density (OD) in microtiter plates containing serial dilutions of fungicides has now been widely implemented (e.g. ^49,10,19^) and works best for fungi that exhibit yeast-like growth. However, a pathogen such as *Z. tritici* tends to produce variable amounts of melanin and can switch from blastospore growth to mycelial growth under stressful conditions, which can affect the accuracy of OD measurements ^37^. For pathogens like *Z. tritici*, a non-toxic redox dye called resazurin (RZ) can provide an alternative measure of growth in microtiter plates ^51, 53^ that is unaffected by changes in growth morphology or melanin production. RZ has a blue color that irreversibly turns pink upon exposure to metabolic activity and currently provides a routine bioassay for measuring cytotoxicity ^7^. RZ in microtiter plates has already been used to measure fungicide efficacy for *Aspergillus spp.* infecting humans and for *Alternaria alternata* infecting crops ^55, 39, 53^. Here we demonstrate the application of RZ to measure fungicide efficacy in *Z. tritici*.

Different measures of fungicide efficacy come with different limitations. An MIC value fluctuates according to the range of fungicide concentrations, the timing and method of collecting data, and the initial spore concentration ^48^. Despite its simplicity and a clear endpoint indicating the presence or absence of growth, an MIC is tied to a single fungicide concentration and does not consider a potentially wide spectrum of dose-response patterns of fungicide efficacy. In contrast, an EC50 (the effective concentration of a fungicide that reduces growth by 50%) that is calculated across several concentrations can provide a more comprehensive insight into fungicide efficacy and allow for a more robust comparison between different studies. EC50 measurements can differentiate between fungistatic (inhibiting growth) and fungicidal (killing fungal cells) effects, unlike MIC ^38^. However, EC50 measures require more extensive data compared to MIC measures, and EC50 calculations are more complex because they follow a non-linear model. Several studies have already used EC50 to measure fungicide efficacy in *Z. tritici* populations (e.g. ^10, 33, 23^). Growth inhibition assays, which lie between MIC and EC50 measures, have also been extensively used to measure reductions in *Z. tritici* growth in fungicide-amended media compared to growth on Petri plates without fungicide ^29^.

A continuing challenge in fungicide resistance research is deciding when a pathogen strain should be called “resistant” and when it should be called “sensitive”. It is relatively easy to distinguish between resistant and sensitive strains when there is an obvious step change between categories, as seen for QoI fungicides ^12^. It is much more challenging to distinguish between categories when resistance is a quantitative trait with a continuous distribution, as found for DMI fungicides ^22^. An additional complication is that different investigators may use different thresholds to define resistant isolates. For example, Harper ^16^ used microtiter plates to estimate the EC50 in *Botrytis cinerea*, followed by a t-test or Mann-Whitney U-test (for normally or non-normally distributed datasets, respectively) to determine a threshold of resistance. Consequently, the EC50 values were classified into low-, medium-, and high-resistance categories based on the tested fungicide and the resulting variation. A similar classification of strains of *Pyricularia graminis-tritici* causing wheat blast was reported by Vicentini et al. (2021), who used the Scott-Knott test (*p*-value = 0.05) to categorize EC50-based phenotypic groups. A promising approach is based on the Silhouette coefficient ^42^ which can statistically define the optimal number of clusters within a phenotypic dataset (e.g., EC50), and provide a more statistically robust identification of resistant strains in pathogen populations ^4^. After defining resistant strains, it is common to study them more intensively to identify specific mutations that encode the resistance and then use these mutations to identify other resistant strains ^23, 33^.

Here we report an extensive investigation of fungicide resistance in a unique collection of *Z. tritici* isolates assembled from a single experimental field in Eschikon, Switzerland. The isolates were sampled in 2016 from a field where 335 elite European winter wheat cultivars were naturally infected by STB despite intensive applications of fungicide mixtures ^25^. By collecting strains at different points in time from known hosts growing in the same field, we could investigate several processes associated with fungicide resistance. In particular, the current work aimed to:

(i) Measure simultaneously the EC50s of six active ingredients in over 1000 *Z. tritici* strains collected from the same field at two time points during a single growing season.
(ii) Quantify the diversity of fungicide phenotypes that exist within fungicide-treated fields and determine if this diversity changes over the course of a growing season.
(iii) Identify isolates that are resistant to individual or multiple active ingredients and quantify changes in frequency of resistant isolates over time.
(iv) Determine whether host genotype affects resistance to different active ingredients, i.e. is there compelling evidence for host-fungicide-pathogen interactions?

## Materials and Methods

### Fungal materials

A replicated field experiment was conducted to evaluate quantitative resistance to STB in 335 elite winter wheat (*Triticum aestivum*) varieties grown under field conditions ^25^. The experiment was conducted in two biological replicates during the 2015-2016 growing season, with the two replicates separated by ∼100 m in the Eschikon field station of ETH Zurich, Switzerland (coordinates 47.449°N, 8.682°E). All wheat cultivars were infected naturally by the local population of *Z. tritici*, with many primary infections likely coming from airborne ascospores originating from other wheat fields growing in the area. All 1.2 x 1.7 m plots were sown on 13 October 2015. The plots were treated three times with different fungicidal mixtures, namely (i) Input®, Bayer (a mix of spiroxamine at 300 g/liter and prothioconazole at 150 g/liter, at a dose of 1.25 liters/ha, at Growth Stage 31 (GS 31, Zadoks, et al., 1974); (ii) Aviator® Xpro, Bayer (a mix of bixafen at 75 g/liter and prothioconazole at 150 g/liter, at a dose of 1.25 liters/ha, GS 51); and (iii) Osiris®, BASF (a mix of epoxiconazole at 56.25 g/liter and metconazole at 41.25 g/liter, at a dose of 2.5 liters/ha, GS 65 that corresponds to anthesis). Sixteen leaves showing symptoms of STB were sampled randomly from each plot at two times during the season (32 leaves in total). The first collection (C1) was sampled on 20 May at GS41 and a third collection (C3) was sampled on 4 July at GS75-85. A second smaller collection (C2) sampled on 17 June was not included in these analyses. Groups of eight leaves were mounted on A4 paper, scanned, and analyzed using ImageJ to measure the percentage of leaf area covered by lesions (PLACL), pycnidia density within lesions (ρlesions), and pycnidia density within leaves (ρleaf ^25^) as quantitative indicators of resistance to STB.

To isolate *Z. tritici* from these leaves, a cirrus from a single pycnidium was plated on a Petri dish containing yeast malt sucrose agar (YMSA) with kanamycin (50 µg.ml^-1^). After incubating at 18°C for 7 days, a single-spore colony was transferred to 35 ml of yeast sucrose broth (YSB) and placed on an orbital shaker (180 rpm) for 7 days at 18°C. A dense concentration of blastospores was preserved on silica gel and stored at -80°C for future use ^47^. To recover the isolates from long-term storage, dried spores were transferred into 20 ml of YSB (10 g sucrose and 10 g yeast extract in 1 liter of ddH_2_O + 50 mg.l^-1^ streptomycin + 50 mg.l^-1^ chloramphenicol). After 7 days of incubation at 180 rpm at 18°C in the dark, the blastospore suspension was filtered through two layers of sterile cheesecloth, then centrifuged at 3500 rpm (2851g) at 4°C for 15 min. The spore pellet was resuspended with sterile dd H_2_O and the blastospore concentration was adjusted to 8x10^5^ spore.ml^-1^ using KOVA counting slides and a light microscope. In total, 410 of the isolates from C1 and 595 of the isolates from C3 were used in the experiments of fungicide efficacy reported here.

### Microtiter plate assays

We tested fungicidal effects in sterile 96 well microtiter plates. Each *Z. tritici* isolate was added at the same concentration (8x10^5^ blastospores.ml^-1^) to a column of eight wells; seven with media containing a fungicide diluted to seven concentrations and one control well containing media but no fungicide. Every isolate was replicated three times in different microtiter plates. Each plate included one control column that did not receive any blastospores (i.e., with no fungal growth). The position of the control column was completely randomized with respect to the other 11 isolate-containing columns on the same plate. Plates were sealed with parafilm, labeled with a unique QR code, and incubated at 18°C for seven days in the dark and at 60% relative humidity to minimize the evaporation of liquid from the plates. The plates were distributed on metal mesh shelves in an incubator (Kälte 3000 AG, 2008, Switzerland) using a completely randomized design. After seven days, we scanned the plates and the associated QR code using a flatbed scanner at 600 dpi.

### Calibration of Resazurin

Resazurin dye (RZ) was dissolved into sterile ddH_2_O and sterilized through a 45 µm filter to a final concentration of 5000 mg.l^-1^. Two *Z. tritici* strains, c1_25_2B2 and c1_290_9H1, were used to conduct this calibration experiment. To determine the optimum concentrations of RZ and blastospores to use in the main experiments, we tested serial dilutions of RZ at different concentrations (350, 300, 250, 200, 180, 150, 100, 80, 50, 40, 30, and 20 mg. l^-1^) against serial dilutions of blastospores at different concentrations (1 x 10^5^, 2 x 10^5^, 4 x 10^5^, 6 x 10^5^, 8 x 10^5^, 10 x 10^5^, 12 x 10^5^, 15 x 10^5^ spores.ml^-1^), providing 96 different combinations of RZ and blastospores in each plate. This optimization experiment was replicated twice for each isolate. Our goal was to determine the lowest concentration of RZ that would detect the greatest range in color change (i.e., from blue to red) as a measure of fungal metabolism that could be used as a proxy of fungal growth. The plates were scanned using a flatbed scanner at different times (1, 24, 48, 72, 96, 120, 144, and 168 hours after adding spores) to measure the amount of red color produced in response to fungal metabolism.

### Measuring fungicide efficacy

We used the RZ as a metabolic indicator of the amount of *Z. tritici* growth in each well based on measuring the induced red color (i.e., resorufin) using digital image analysis. The serial dilution of each fungicide was adjusted to fill one column in the microtiter plate with 8 different concentrations. Five of the tested active ingredients (AIs, prothioconazole, metconazole, epoxiconazole, spiroxamine, and bixafen) were applied to the field experiment where the *Z. tritici* isolates originated. We added propiconazole as a 6^th^ AI that has been applied for many years in European wheat fields. The tested fungicide ranges were chosen to enable some growth of fungal isolates at low concentrations while aiming to achieve no growth or minimal growth at the highest concentration. We used the following serial dilutions: 0, 3.75, 7.5, 18.75, 37.5, 112.5, 150, and 225 mg.l^-1^ for propiconazole and prothioconazole; 0, 3.75, 7.5, 18.75, 37.5, 75, 112.5, and 150 mg.l^-1^ for metconazole; 0, 0.75, 3.75, 7.5, 18.75, 37.5, 75, and 187.5 mg.l^-1^ for epoxiconazole; 0, 0.08, 0.38, 0.75, 1.5, 3.75, 7.5, and 75 mg.l^-1^ for spiroxamine; and 0, 0.008, 0.075, 0.375, 0.75, 1.5, 7.5, and 75 mg.l^-1^ for bixafen.

After analyzing the complete dataset for all isolates, we identified subsets of isolates (52, 53, 88, 70, 29, and 3 isolates for propiconazole, prothioconazole, metconazole, epoxiconazole, spiroxamine, and bixafen, respectively) that required either a higher or lower range of fungicide concentrations to establish their EC50 values. We re-tested those isolates using a different range of serial dilutions as described in Supplemental Table 1.

### Analyzing the digital images

We created a series of shell and ImageJ Macro (IJM) scripts to create and read the QR codes, sort the images according to the plate name (i.e., the active ingredient, replicate, and date of testing), and analyze the red color intensity in each well of each microtiter plate (Supplemental Files 1, 2, and 3). The red color intensity was used as a proxy for the fungal biomass in each well. We modified an existing macro for reading red color intensity in microtiter plates ^1^ ( Supplemental File 3). Each well in a plate was identified and labeled based on its position in the plate. Then each well image was analyzed to separate its color components (RGB; Red, Green, and Blue) and the red intensity of each pixel was calculated on a scale of 0-255 using ImageJ ^28, 1^. The degree of redness mirrors the intensity of the metabolic activity of the fungus (i.e., the higher its metabolism, the more intense the red color) and the metabolic activity was used as a proxy of the total fungal biomass present in each well. Based on the well position in the plate, we could analyze red intensity values as a function of fungicide concentration.

### Calculating EC50 values

The EC50 value was calculated based on the growth of each isolate over eight concentrations by applying a nonlinear model with four parameters using the following formula:

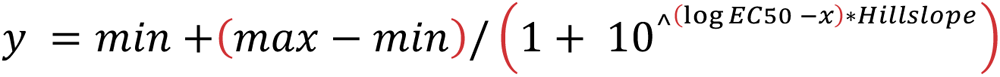

Where *y* = the isolate response to the active ingredient (i.e., measured by the redness value as an indicator of fungal growth), *min* = minimum red value within the active ingredient spectrum of the isolate under investigation (at the highest AI concentration), *max* = maximum red value within the active ingredient spectrum of the isolate under investigation (at the zero AI concentration), EC50 is the concentration of active ingredient that can suppress the metabolic activity (i.e., growth) of a fungal isolate by 50%, and *x* = the concentration of active ingredient corresponding to *y* fungal growth. The individual red values of each replicate were integrated into the model, hence the average value of each data point (i.e., AI concentration) was used to generate the EC50 curve and the fitted EC50 value.

### Other statistical analyses

The optimization of experimental conditions was analyzed using SAS software ^45^. ImageJ ^46^ and command line scripts (Supplemental files 1, 2, 3) were used to generate the raw data used to calculate EC50 values for each isolate. SPSS software ^20^ was used to measure the correlation between EC50 replications. Python-Pandas ^36^ was used to manipulate the data and define different clusters of resistant isolates based on population mean values and their standard deviations. The remainder of the analyses and plotting of results was performed with R software ^40^ and R Studio ^43^. The EC50 values were normalized (*zi*=*xi*-min(x)/max(x)-min(x)) because the scales varied for each AI (e.g. for bixafen values were <10, while for propiconazole values ranged up to 161) and log transformed to meet the assumptions of linear models. To estimate broad sense heritability (*H^2^*) of EC50, we used a linear mixed effect (LME) model and variance partitioning. For the LME, the log transformed and normalized EC50 value was the response variable, collection time was the fitted factor, and random factors were host, AI, replicate and isolate. Variance components were extracted and used to calculate *H^2^*. To analyze each AI separately, two approaches were used: the first used the value for each technical replicate as the response variable and included replicate in the model as an explanatory variable (y^∼^ collection + replicate + host); the second used the mean value across the three technical replicates as the response variable (y^∼^ collection + host). Although the first approach is less conservative, it takes into account possible microenvironment effects during incubation and may reveal more subtle effects because of the 3X larger number of observations. After fitting linear models and performing an analysis of variance, we performed multiple pairwise comparisons using least significant differences (LSD) and adjusted the *p*-values for multiple testing using the Bonferroni correction. To investigate the effect of collection time, host and the interaction between AI and host, the mean EC50 across three replicates was used as the response variable, while collection time, host, AI and the interaction between host and AI were explanatory variables. We explored multiple approaches to classify isolates as resistant or sensitive to each AI as described in detail in the Results section. Using the most conservative classification scheme, we investigated how the frequency of resistance to each AI was influenced by collection time and host using a generalised linear model with a binomial distribution. To investigate how host was related to an isolate’s resistance to different combination of AIs, binomial tests were used to test if the probability of multidrug (and combinations of dual-, triple-, quadruple- or quintuple-drug) resistance was higher than expected based on multiplying the individual probabilities of each AI separately. R software was also used to calculate the Scott-Knott and K-means thresholds of fungicide resistance in order to cluster the isolates into different categories of resistance.

## Results

### Developing a digital image analysis pipeline

The microtiter plates with RZ showed different gradients of colors which reflected the cumulative growth of each isolate across the serial dilutions of each active ingredient. The uninoculated control column in each plate remained blue and did not show any change in color across all eight concentrations over the entire experiment.

We tested 12 different concentrations of RZ in the absence of fungicides and measured the red color scores across 1 to 168 hours of incubation. We found significant variation among the RZ concentrations for both tested isolates (Figure 1). A concentration between 180-200 mg.l^-1^ was chosen as the lowest RZ concentration that covered a significant fraction of the red color spectrum. Color densities (i.e., gradual changes from blue to red relative to varying concentrations of RZ and blastospores) also varied over time. More than one peak can be seen between the 1 and 120 h post-inoculation periods (Figure 2). Although we found a single peak at 144 h, a more normal distribution was exhibited by the peak seen at 168 h post-inoculation (Figure 2). Based on these findings, we chose 168 h after inoculation as an optimal time point for collecting the data. ANOVA revealed significant variation among blastospore concentrations and RZ concentrations, but no significant differences were found between replications (Table 1). An LSD test of the blastospore concentrations indicated three significantly different categories, leading us to choose the intermediate group (8 x 10^5^ – 12 x 10^5^ spore.ml^-1^) (Supplemental Figure 1) for the main experiment. In summary, these findings led us to choose 168 hours of incubation, 180 ml.l^-1^ of RZ, and 8 x 10^5^ blastospores/ml as the optimum conditions to measure fungicide sensitivity.

**Figure 1.**
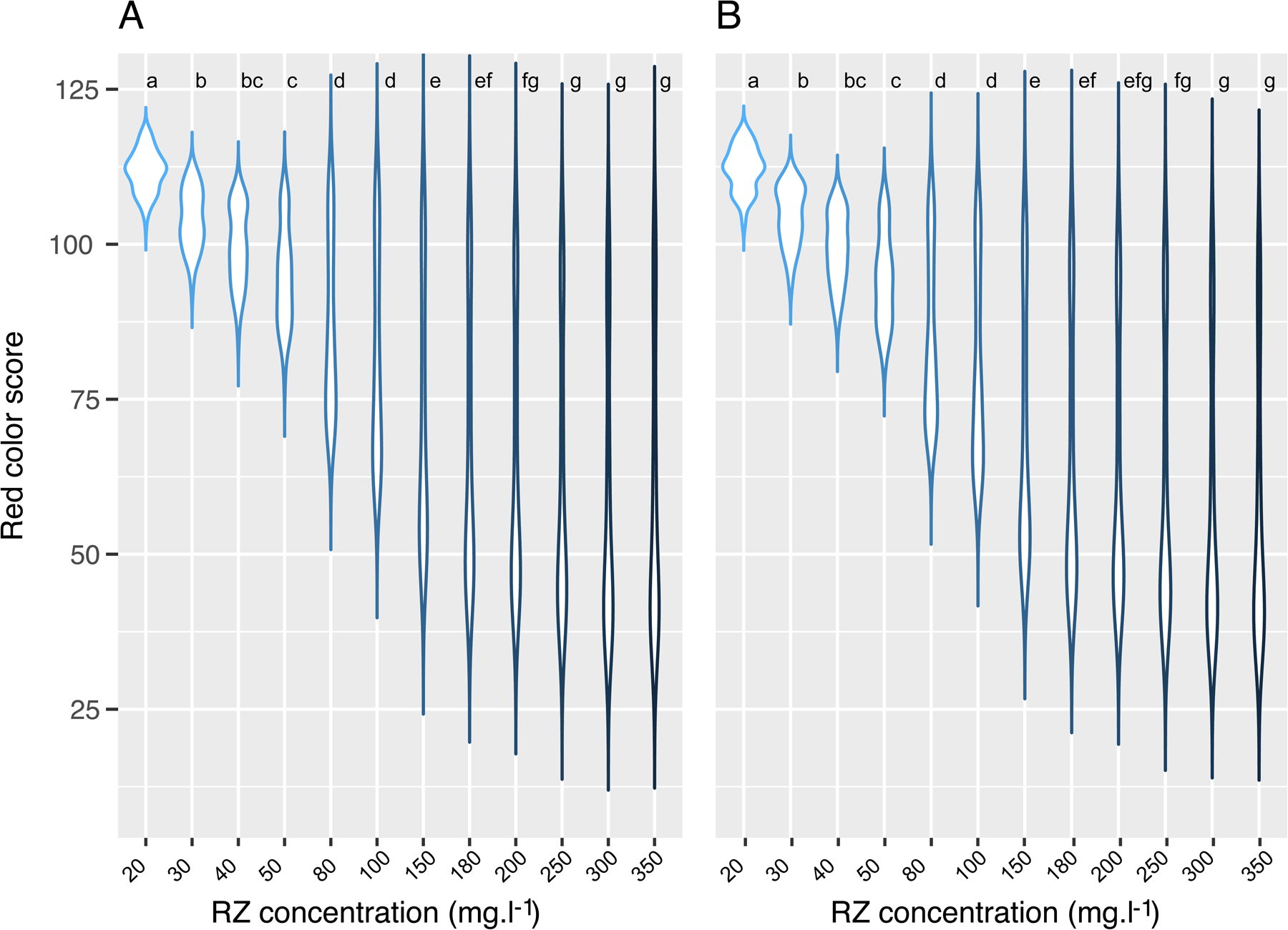
Optimizing resazurin (RZ) concentration through a factorial experiment using two isolates of *Zymoseptoria tritici*; A: c1_25_2B2 and B: c1_290_9H1, each with two repetitions. After applying a Bonferroni correction, the Least Significant Difference (LSD) was used to determine differences among treatments, indicated with letters at the top of each tested RZ concentration. The concentration range of 180-200 mg/l was identified as the minimum level of RZ that captured a significant proportion of the red color spectrum, indicative of fungal metabolic activity.

**Figure 2.**
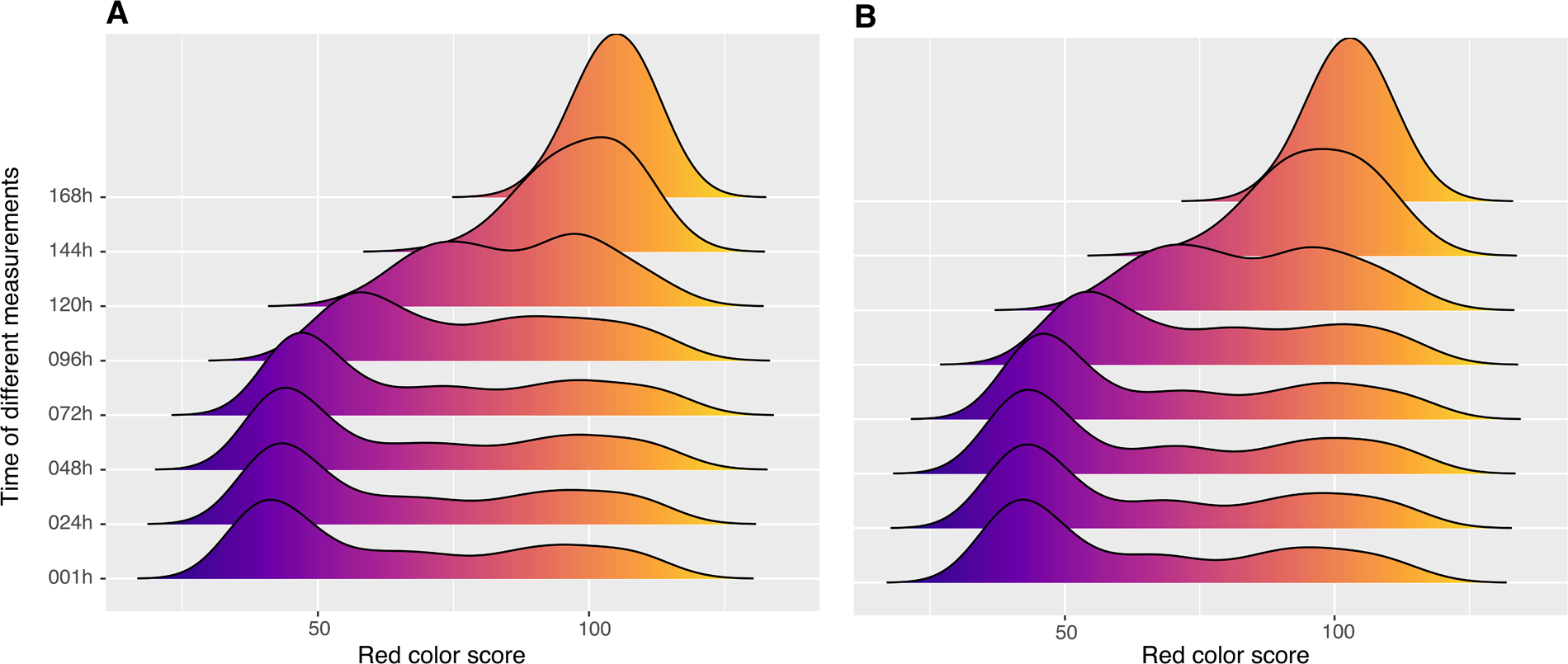
Optimizing the incubation time for data collection in a factorial experiment involving two strains of *Zymoseptoria tritici*; A: c1_25_2B2 and B: c1_290_9H1, each with two repetitions. The experiment, conducted in 96-well microtiter plates, included a horizontal gradient composed of serial dilutions of resazurin and a vertical gradient composed of serial dilutions of blastospores. Red color intensity was measured at eight different time points for each microtiter plate. Each time point includes the complete distribution of red color scores across all 96 combinations of resazurin and blastospore concentrations. The mean values of both replicates were plotted on the x-axis, revealing quantitative, gradual changes from blue to red. A normal distribution was observed at the optimal incubation time of 168 hours.

**Table 1.** Analysis of variance for effects of resazurin (RZ) concentration and spore concentration on red color intensity in a microtiter plate assay of fungal metabolic activity.

| Source | Df | Sum Sq | Mean Sq | F value | Pr(>F) |
| --- | --- | --- | --- | --- | --- |
| Isolate | 1 | 1239 | 1239 | 4.55 | 0.033 * |
| Spore concentration | 7 | 25208 | 3601 | 13.226 | <2e-16 *** |
| RZ concentration | 11 | 1064252 | 96750 | 355.341 | <2e-16 *** |
| Replication | 1 | 1 | 1 | 0.004 | 0.948 |
| Residuals | 3051 | 830709 | 272 |  |  |

### Measuring fungicide sensitivity

The replications of the same isolate showed the same degree of red color intensity over different fungicide concentrations (Supplemental Figure 2). EC50 values of biological replicates were compared to check the reproducibility of the method. Linear regression was calculated using a Bayesian model in SPSS ^20^. The resulting R^2^ ranged from 0.98 to 0.99 (*p*-value <0.05) for each tested AI. Significant pairwise correlations (Pearson *p*-value < 0.001) were found between EC50 datasets for all AIs (Table 2), but the highest correlations were found among propiconazole, epoxiconazole and metconazole, suggesting that cross-resistance may occur more readily among these DMIs. Cross-resistance between different DMI active ingredients was already reported in *Z. tritici* ^50^ and *Alternaria alternata* ^21^. Cross-resistance was also reported earlier between SDHI and azole fungicides in *Z. tritici* ^26^, in line with our results. The EC50 for each fungicide was calculated separately for the C1 and C3 collections of isolates, providing an average across replicates for 410 and 595 isolates, respectively. The mean (and range) EC50 in mg.l^-1^ were: 0.39 (0.01 – 6.53) for bixafen; 24.7 (0.01 – 127.60) for epoxiconazole; 17.99 (0.07 – 118.20) for metconazole; 26.79 (0.42 – 161.30) for propiconazole; 18.29 (0.01 – 99.88) for prothioconazole; and 4.34 (0.48 – 25.46) for spiroxamine (Table 3). The variance component analysis, using a linear mixed effects model, identified a significant broad-sense heritability (*H^2^*=0.53) associated with the EC50 values (Supplemental Table 2). A similar range of EC50 values was reported for bixafen by several researchers ^9, 41^. However, Birr ^5^ reported that the EC50 of propiconazole did not increase between 1999 and 2020 while also reporting a continuing decrease in sensitivity to prothioconazole, with a maximum EC50 value of < 10 mg.l^-1^. Torriani ^50^ reported EC50 values for prothioconazole ranging from 0.01-100 mg.l^-1^ which is in line with our results. Similarly, Heick ^18^, reported a range from 0.01-100 mg.l^-1^ for prothioconazole in the Nordic-Baltic region. In contrast, the EC50 of epoxiconazole ranged from 0.01-10 mg.l^-1 18^. However, in another report, one isolate had an EC50 value > 100 mg.l^-1^ for epoxiconazole ^19^. In summary, the EC50 values we recorded using RZ dye largely overlapped with the values reported by other labs.

**Table 2.** Correlation matrix between fitted EC50 datasets for six active ingredients. The upper diagonal shows the correlation coefficients and the lower diagonal shows the corresponding *p*-values.

|  | EC50 Bixafen | EC50 Epoxiconazole | EC50 Metconazole | EC50 Propiconazole | EC50 Prothioconazole | EC50 Spiroxamine |
| --- | --- | --- | --- | --- | --- | --- |
| EC50 Bixafen |  | 0.55 | 0.54 | 0.70 | 0.15 | 0.47 |
| EC50 Epoxiconazole | 1.11E-71 |  | 0.70 | 0.70 | 0.23 | 0.48 |
| EC50 Metconazole | 9.27E-69 | 1.46E-132 |  | 0.73 | 0.29 | 0.49 |
| EC50 Propiconazole | 3.30E-61 | 5.84E-136 | 7.33E-155 |  | 0.26 | 0.52 |
| EC50 Prothioconazole | 6.01E-06 | 3.00E-12 | 5.73E-19 | 1.93E-15 |  | 0.16 |
| EC50 Spiroxamine | 2.03E-50 | 2.19E-53 | 2.69E-55 | 4.24E-65 | 6.31E-07 |  |

**Table 3.** Minimum, maximum and mean EC50 values for six active ingredients for the C1 and C3 collections individually and combined together.

| Active ingredient | Collection | Bixafen | Epoxiconazole | Metconazole | Propiconazole | Prothioconazole | Spiroxamine |
| --- | --- | --- | --- | --- | --- | --- | --- |
| Minimum | C1 | 0.01 | 0.19 | 0.07 | 0.42 | 0.23 | 0.48 |
|  | C3 | 0.01 | 0.01 | 0.11 | 0.51 | 0.01 | 0.58 |
| Maximum | C1 | 6.53 | 127.60 | 118.20 | 161.30 | 99.88 | 25.46 |
|  | C3 | 2.00 | 127.20 | 111.90 | 151.70 | 89.59 | 21.04 |
| Mean | C1 | 0.52 | 34.50 | 22.19 | 31.29 | 18.44 | 4.62 |
|  | C3 | 0.28 | 17.99 | 14.97 | 23.59 | 18.18 | 4.15 |
| Mean | C1 & C3 | 0.39 | 24.70 | 17.99 | 26.79 | 18.29 | 4.34 |
| Standard deviation | C1 & C3 | 0.46 | 26.78 | 18.27 | 26.35 | 12.72 | 2.94 |

### Categorizing isolates based on their fungicide sensitivity

We compared several approaches to identify resistant isolates for each fungicide. The first approach, used as a control, treated the mean EC50 value of the total dataset for each AI as a threshold, with all EC50 values above the mean value indicating resistant isolates. Using this approach, we counted 350 (35%) isolates resistant to bixafen, 338 (34%) isolates resistant to epoxiconazole, 327 (32%) isolates resistant to metconazole, 349 (35%) isolates resistant to prothioconazole, and 351 (35%) isolates resistant to spiroxamine (Table 4). The second approach was much more stringent, defining an isolate as resistant if its EC50 is at least one standard deviation above the population mean value (i.e., mean + SD). Using this approach, we counted 77 (8%) isolates resistant to bixafen, 155 (15%) isolates resistant to epoxiconazole; 131 (13%) isolates resistant to metconazole; 123 (12%) isolates resistant to propiconazole; 87 (9%) isolates resistant to prothioconazole, and 122 (12%) isolates resistant to spiroxamine (Table 4).

**Table 4.**
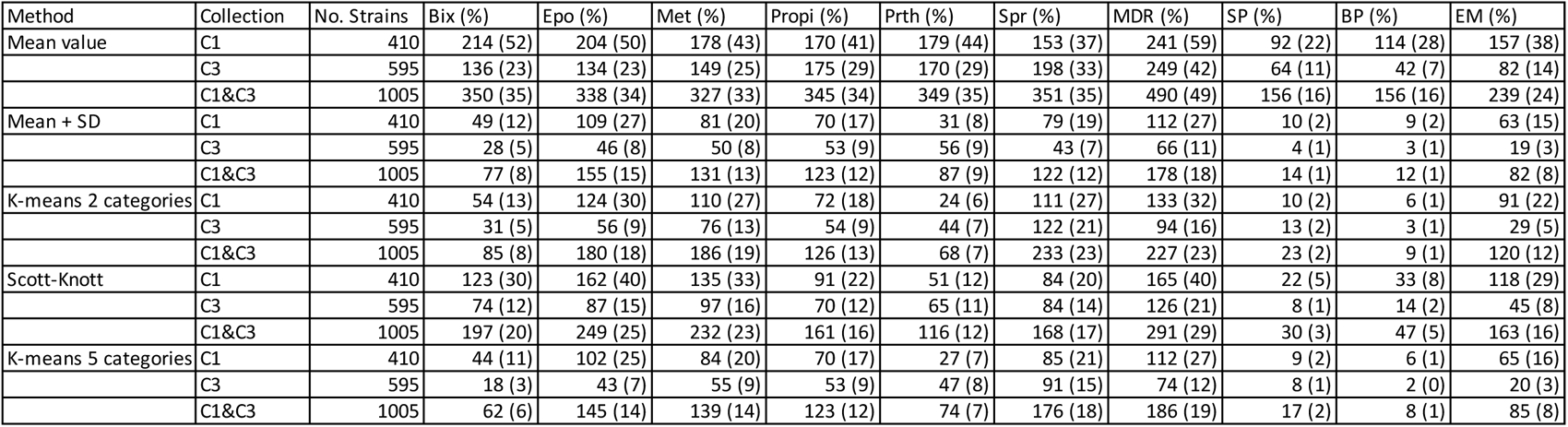
Numbers and frequencies of isolates defined as resistant to each of six active ingredients, and different combinations of active ingredients, based on five different categorical methods. The C1 and C3 collections are analyzed individually and combined together. Differences among the five different categorical methods are described in the main text. Bix = bixafen. Epo = epoxiconazole. Met = metconazole. Propi = propiconazole. Prth = prothioconazole. Spr = spiroxamine. MDR includes all isolates that were resistant to at least two active ingredients. Dual-drug resistances include SP (Spr + Prth), BP (Bix + Prth), and EM (Epo + Met).

In the third approach, we sought to classify the isolates into five discrete categories, namely highly resistant (HR), resistant (R), moderately resistant (M), sensitive (S), and highly sensitive (HS) by applying the Scott-Knott test (*p*=0.05) based on ANOVA, an approach that was already applied to categorize fungicide resistance ^54^. This approach identified 21, 22, 24, 22, 19, and 17 distinguishable categories for bixafen, epoxiconazole, metconazole, propiconazole, prothioconazole, and spiroxamine, respectively. Since it is ANOVA-based, this method has the disadvantage that it analyzes the EC50 values of the replicates separately and not the fitted EC50 values that integrate all replicates. We assigned the resulting groups into five resistance categories (i.e., HR, R, M, S, and HS) by using the EC50 density distribution (= frequency/class width) (Figure 3) and taking into consideration the visible peaks in the distribution that we treated as representing different classes of resistant isolates. After combining the categories R and HR to define resistant isolates, we counted 197 (20%), 249 (25%), 232 (23%), 161 (16%), 116 (12%), and 168 (17%) resistant isolates for bixafen, epoxiconazole, metconazole, propiconazole, prothioconazole, and spiroxamine respectively.

**Figure 3.**
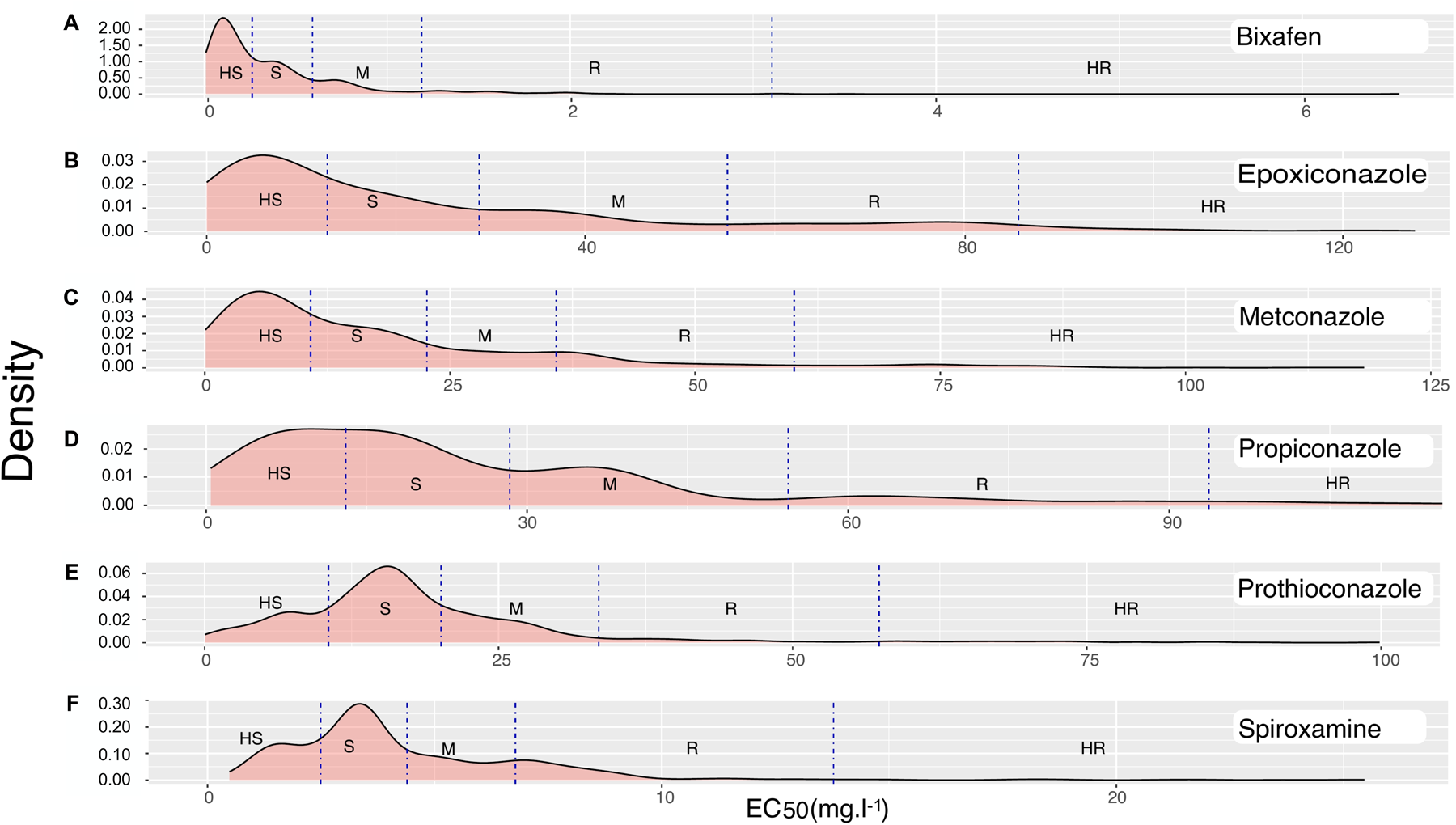
Density distribution of EC50 values for the tested active ingredients, with blue vertical lines delineating borders between different resistance categories based on K-means clustering with five categories. The datasets include bixafen (A), epoxiconazole (B), metconazole (C), propiconazole (D), prothioconazole (E), and spiroxamine (F). Resistance categories are denoted as HS (highly sensitive), S (sensitive), M (moderate resistance), R (resistant), and HR (highly resistant). The horizontal axes represent the range of EC50 values for the active ingredients in mg/l. Isolates in the HR and R categories were categorized as resistant while all other isolates were categorized as sensitive.

The fourth approach used *K*-means clustering based on Euclidean distances ^32, 3, 4^. The silhouette coefficient was calculated for *K* values ranging from 2 to 15 and then plotted against the corresponding *K* value (Supplemental Figure 3). The optimal number of groups in our datasets following the silhouette method was two, which split the data for all fungicides into two categories (Supplemental Figure 3). An advantage of the *K*-means approach is that it can be used to identify optimal subgroups based on a user-defined number of categories. For the fifth approach, we used *K*-means clustering to define five categories (HR, R, M, S, and HS, Figure 3), and we called an isolate resistant if it was classified into the HR or R categories. This analysis identified 62 (6%) isolates resistant to bixafen, 145 (14%) resistant to epoxiconazole, 139 (14%) resistant to metconazole, 123 (12%) resistant to propiconazole, 74 (7%) resistant to prothioconazole, and 176 (18%) resistant to spiroxamine.

We compared the identification of resistant isolates based on mean EC50 values, mean + SD, Scott-Knott, *K*-means (*K*=2) and *K*-means (*K*=5). The mean value approach used as a control identified the highest number of resistant isolates for each AI, while the Scott-Knott method typically identified the second highest number of resistant isolates. The other three approaches found lower numbers of resistant isolates that in most cases did not differ substantially from each other (Table 4), with the numbers between the different approaches being highly correlated (R^2^=0.86, *p*-value = 0.006). We decided to use the *K*-means (*K*=5) identification of resistant isolates for all further analyses because this approach offered more flexibility in separating different categories of isolates while also being sufficiently conservative in identifying resistant isolates.

Isolates exhibiting multidrug resistance, using the *K*-means (*K*=5) definition of resistance, were categorized as (i) Dual-drug resistant (DDR), based on the fungicide combinations that were applied to the field experiment, resulting in 17 (1.7%) isolates resistant to SP (spiroxamine + prothioconazole), 8 (0.8%) isolates resistant to BP (bixafen + prothioconazole), and 85 (8.3%) isolates resistant to EM (epoxiconazole + metconazole), and (ii) Multidrug resistant (MDR) based on resistance to any combination of fungicides including at least two AIs, which resulted in 186 (18.9%) isolates exhibiting MDR. The EM-resistant isolates occurred at the highest frequency (8.3%).

### The effects of collection time on EC50 and single- and multidrug resistance

The EC50s were significantly higher in the first collection (Supplemental Table 3, Figure 4), with the same pattern found across most AIs. When using the EC50 for each replicate, the EC50 was significantly higher in the first collection for all AIs except spiroxamine (Supplemental Table 3). When using mean EC50 across replicates, the effect of collection time was significant for all but two AIs: spiroxamine and prothioconazole (Supplemental Table 4). The frequency of single drug resistance was significantly higher for isolates collected during the first collection with one exception, prothioconazole (Supplemental Table 5). Overall MDR and DDR to EM and BP were higher during the first collection (Supplemental Table 5, Figure 4). For the rest of the analyses, we include collection time in our models to control for these differences.

**Figure 4.**
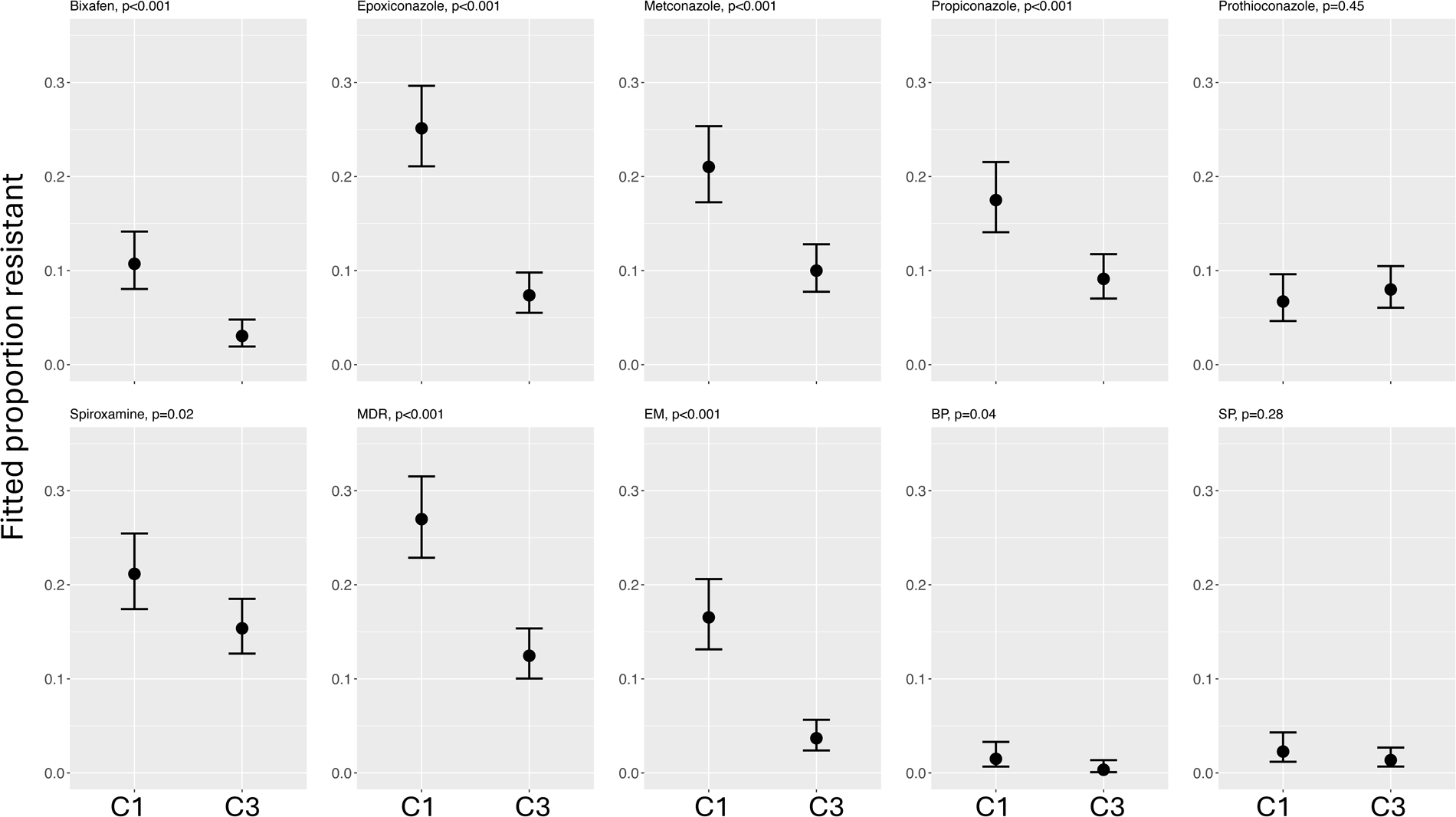
Fitted means (±95% confidence intervals) of the proportion of resistant isolates for each active ingredient, and dual- and multidrug resistant combinations in the first field collection (C1) and the third field collection (C3). Dual-drug resistance combinations are: BP (bixafen and prothioconazole), SP (spiroxamine and prothioconazole), and EM (epoxiconazole and metconazole). Multidrug resistance (MDR) is resistance to any combination of the six tested active ingredients. Significance is indicated with *p*-values in each panel.

### The effects of Host-Fungicide-Pathogen (H-F-P) interactions on EC50

The *Z. tritici* isolations were made from 17 host genotypes that differed in their resistance to STB. We excluded the cultivars Garcia and Florett from this analysis because only two and five isolates, respectively, came from these cultivars, leaving 15 cultivars for comparisons. A significant H-F-P interaction was found when data with all AIs was combined: host, AI and their interaction explained a significant amount of the variation in the EC50 of each isolate (Supplemental Table 6). When considering each AI separately and each replicate as an observation, we found that EC50 varied with Host for all AIs (Supplemental Table 3, Figure 5). Individual cultivars hosted isolates with significantly higher EC50s for an AI compared to other cultivars; for example, isolates from Zinal showed the highest EC50 for prothioconazole, while isolates from Cassiopeia showed the highest EC50 for both bixafen and metconazole (Figure 5). If we use the mean across replicates, fewer significant host differences were observed; host did not explain variation in EC50 of propiconazole, and for epoxiconazole the effect was almost significant (0.056) (Supplemental Table 4, Supplemental Figure 4).

**Figure 5.**
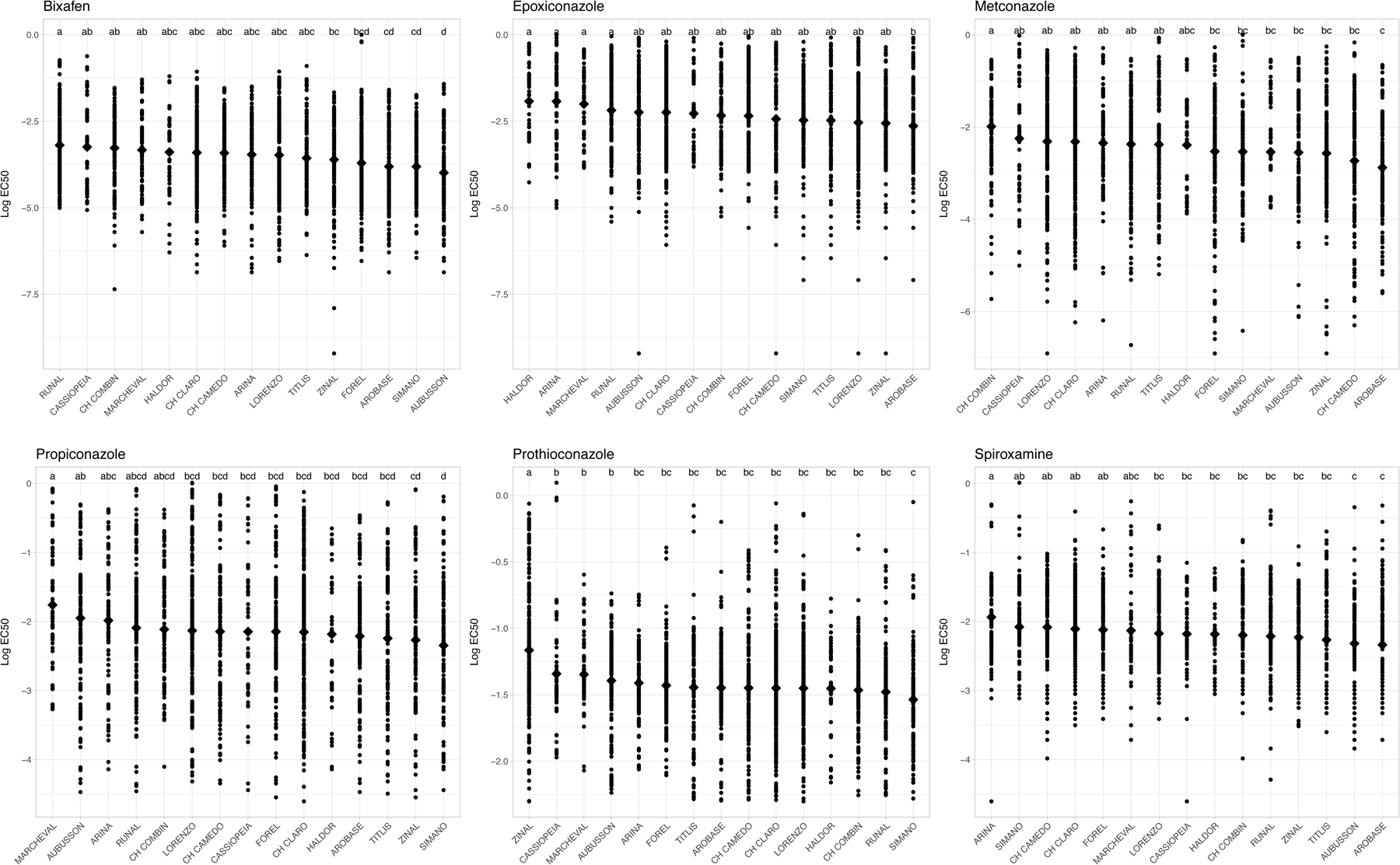
Log transformed and normalized EC50 values (points) for isolates sampled from different cultivars, sorted from highest mean EC50 (diamonds) to the lowest mean EC50. Letters atop each bar (i.e. cultivar) identify significantly different groups. Each panel includes data from the indicated fungicide.

Next, we tested for associations between the average EC50 for an AI on each cultivar with the average degree of STB resistance measured on that cultivar for three different measures of resistance: PLACL, ρlesions and ρleaf, using the data reported in Karisto ^25^. The STB resistance datasets were generated using the same leaves which were the source of the *Z. tritici* isolates described in this paper. We found negative correlations between the average EC50 values and the average degree of STB susceptibility measured for each cultivar for 14 of the 18 comparisons (Table 5, for all pairwise comparisons see Supplemental Table 7), but only three of these were statistically significant. We also observed significant negative correlations between frequency of MDR and ρlesions (-0.82) and ρleaf (-0.60) (Table 5). We note here that higher values for PLACL and pycnidia density reflect higher susceptibility to STB, while higher values for EC50 reflect higher resistance to the associated AI. Hence, a negative correlation between these factors indicates a positive relationship between STB resistance and resistance to the corresponding fungicide.

**Table 5.**
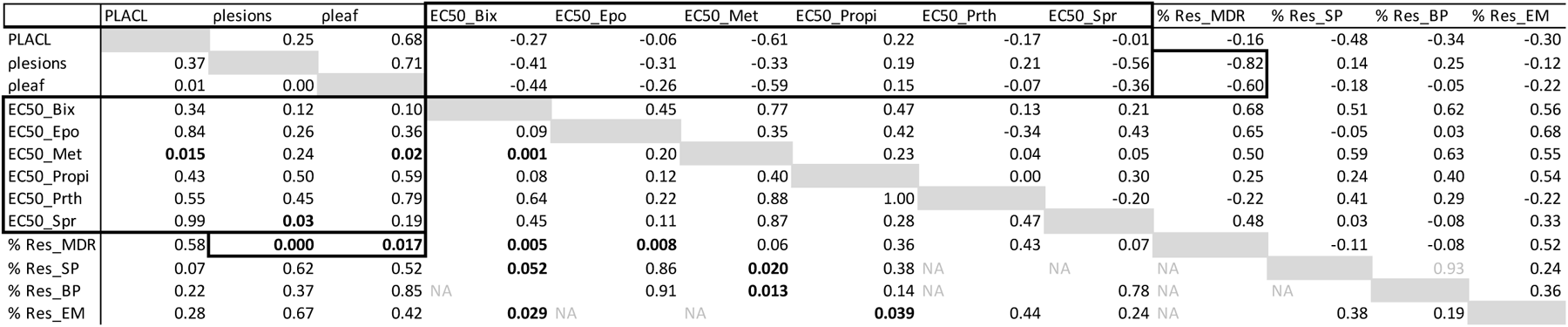
Correlations between EC50 cultivar based (i.e., mean EC50 value of isolates hosted by a certain cultivar), *Zymoseptoria tritici* pathogenicity (PLACAL, pycnidia density, and *ρ*-leaf) and the percentage of Dual- and Multidrug resistant isolates per cultivars. Correlation coefficients are in the upper diagonal and corresponding p-values are in the lower diagonal. *p*-values are only provided for strictly independent variables.

Finally, we compared the average frequency of resistant isolates found for each AI on each cultivar with the average degree of STB resistance exhibited by that cultivar. After controlling for differences between collection times, we found significant differences in the percentage of resistant isolates found on different cultivars for prothioconazole (*F* =60.3, df=14, *p*<0.001) and spiroxamine (*F* =37.3, df=14, *p*<0.001) (Figure 6), but not the other AIs (Supplemental Table 5). Arina hosted the *Z. tritici* population with the highest EC50 for spiroxamine and it also had the highest frequency of isolates that were scored as resistant to spiroxamine. The same pattern was found for Zinal and prothioconazole. As before, the correlations between susceptibility to STB and frequency of resistance to an AI or a combination of AIs were mostly negative (23 out of 30 comparisons), but only three of them were significant (Supplemental Table 7). The highest correlation (-0.82) was found between pycnidia density within lesions (ρlesions) and the frequency of isolates showing MDR (Table 5).

**Figure 6.**
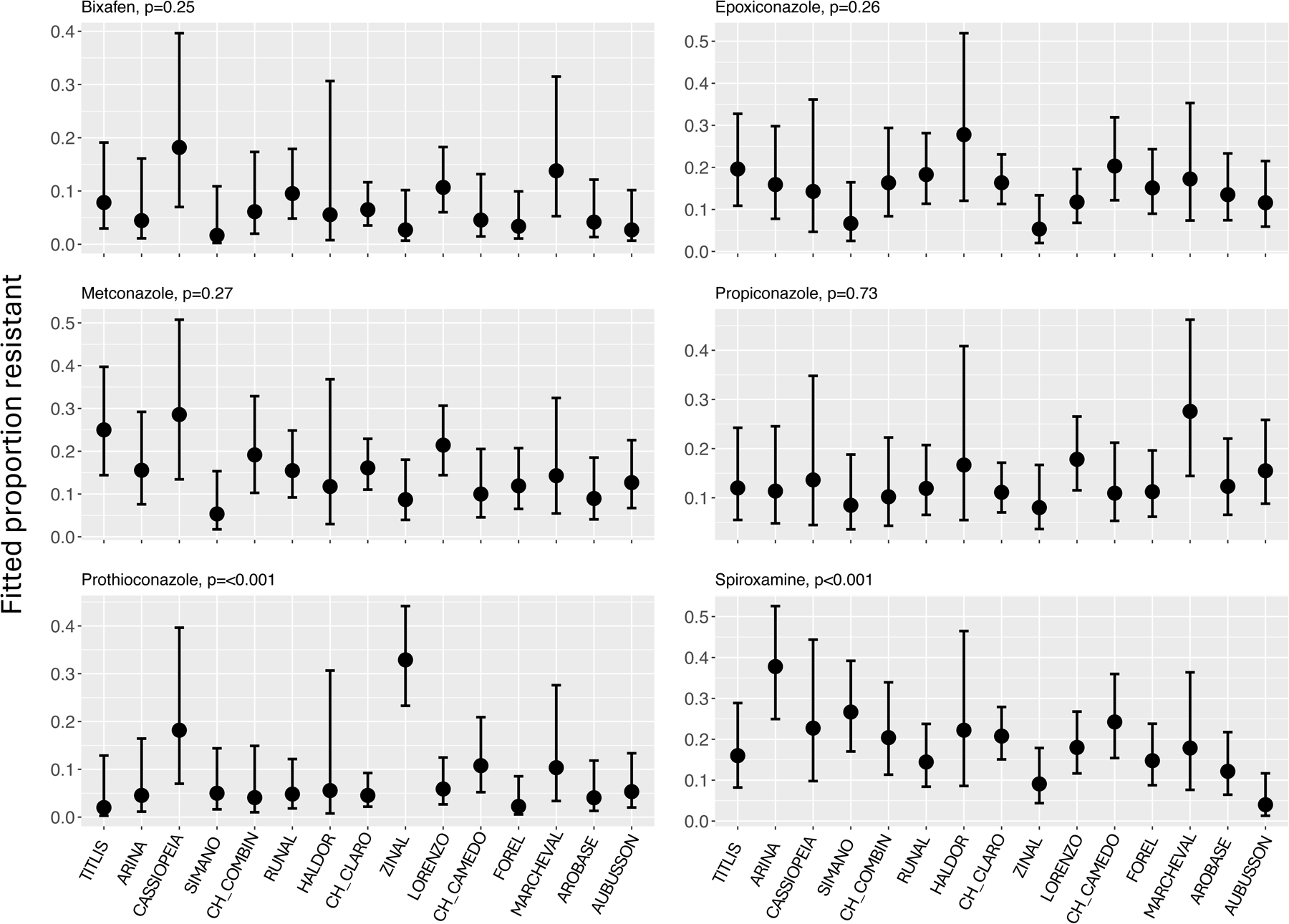
Fitted means (±95% confidence intervals) of the proportion of resistant isolates found for each active ingredient on different wheat cultivars. The cultivars are arranged from highest resistance to septoria tritici blotch (left) to least resistant (right) based on pycnidia density. Significance is indicated with p-values in each panel.

### Multidrug resistance

The probability that isolates have MDR was significantly higher than the null expectation based on multiplying their individual probabilities for nearly all AI combinations (Figure 7, for statistics see Supplemental Table 8). The exceptions were the following dual-drug resistance (DDR) combinations: spiroxamine + prothioconazole (SP), bixafen + prothioconazole (BP), propiconazole + prothioconazole (RP), epoxiconazole + prothioconazole (EP) (Figure 7). We found that MDR varied during the growing season, but did not vary between cultivars (Supplemental Table 5). The C3 collection had a lower proportion of isolates with MDR than the C1 collection (*z*=-5.6, *p<*0.001).

**Figure 7.**
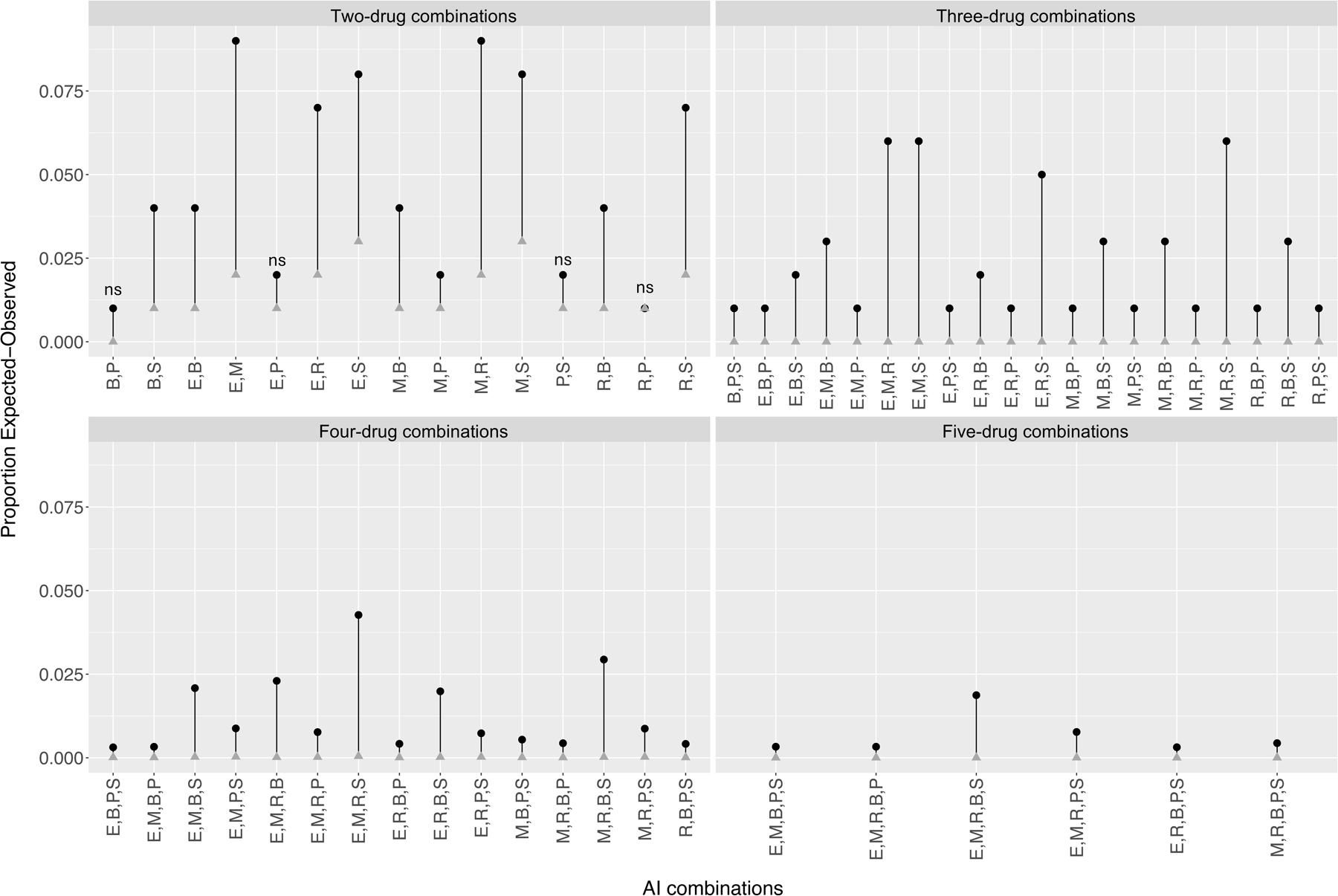
A comparison between the expected (gray triangles) and observed (black dots) proportions of multidrug resistant *Zymoseptoria tritici* isolates within the entire collection of 1005 isolates. Tested active ingredients included bixafen (B), epoxiconazole (E), metconazole (M), prothioconazole (P), propiconazole (R), and spiroxamine (S). All observed proportions were significantly greater than expected except SP, BP, RP, and EP, indicated with ’ns’.

Focusing on the three most common active ingredient combinations applied in the experiment (SP, BP, and EM), we observed differences in DDR among cultivars, ranging from 9-23% of the isolates. EM resistance was the most common dual-drug resistance (Figure 8), but this did not differ amongst cultivars and SP resistance also did not differ amongst cultivars, but the frequency of BP resistance did vary across cultivars (Supplemental Table 5, Figure 8). Cassiopeia stood out for having more than 10% of isolates resistant to SP and BP and was significantly higher than most other cultivars (Figure 8).

**Figure 8.**
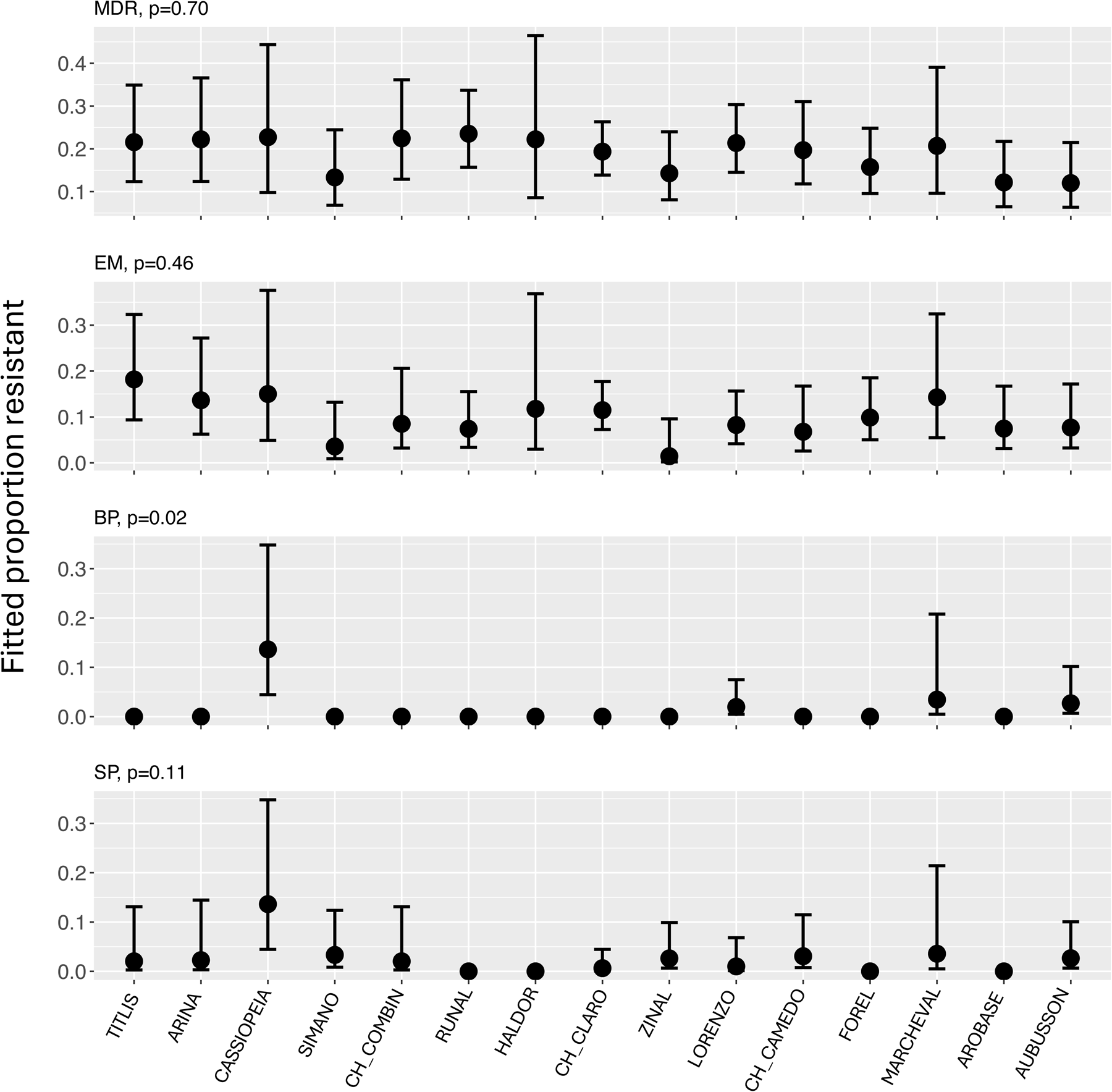
Fitted means (±95% confidence intervals) of the proportion of dual- and multidrug resistant isolates found on different wheat cultivars. Dual-drug resistance combinations are: BP (bixafen and prothioconazole), SP (spiroxamine and prothioconazole), and EM (epoxiconazole and metconazole). Multidrug resistance (MDR) is resistance to any combination of the six tested active ingredients. The cultivars are arranged from highest resistance to septoria tritici blotch (left) to least resistant (right) based on pycnidia density. Significance is indicated with *p*-values in each panel.

## Discussion

*Z. tritici* is already known to adapt rapidly to its local environment by becoming virulent on genetically resistant cultivars ^6, 26^ and resistant to applied fungicides ^22^. In this study, we aimed to characterize the evolution of fungicide resistance during a single growing season in a single naturally-infected field planted with a diverse collection of elite winter wheat cultivars. This field was treated three times during the growing season with five different active ingredients representing the SDHI, DMI, and morpholine modes of action. We found evidence for significant levels of resistance against individual active ingredients as well as high levels of multidrug resistance. We also found strong evidence for significant host-fungicide-pathogen interactions.

### Resazurin dye in microtiter plates provides a new method for measuring EC50 values in *Z. tritici*

Under stress, *Z. tritici* strains often increase melanin production ^29^ and shift from blastospore growth to hyphal growth ^44^. Both processes can affect growth measurements based on the optical density of solutions of fungal spores, so we developed a different method that measures overall metabolic activity as a proxy for fungal growth. Fungicide sensitivity in microtiter plates was based on the RZ color conversion from blue to red associated with total metabolic activity. This method provides quantitative EC50 measurements that are less likely to be affected by the type of growth (i.e., blastospores vs hyphal growth) or the production of melanin under stress. The method is semi-automated and less prone to human error and has the advantage that the analyzed images can be archived for future analyses. The measured EC50 values were in broad agreement with EC50 values reported by other labs using more traditional methods.

### A high diversity of fungicide sensitivities was maintained within a field population over time

The natural field populations of *Z. tritici* sampled in this experiment maintained very high diversity for fungicide sensitivity phenotypes, even after three cycles of selection due to fungicide applications. Despite the strong fungicidal selection, there was no evidence for increasing levels of fungicide resistance over time (Figure 4). The EC50 values ranged over 2-4 orders of magnitude across the >1000 isolates sampled from this naturally infected field. The range of EC50 values found in this single field during a single growing season is similar to the range in EC50 values found for the same fungicides at 55 field trial sites distributed across Europe over a 4-year period ^22^. This agrees with the fine spatial scale over which phenotypic diversity is distributed within field populations of *Z. tritici*, as described in earlier publications^58, 11^. This very high diversity is expected in natural populations not exposed to fungicide selection ^35^, but it was much higher than we expected to find in a field population exposed to three separate fungicide sprays composed of five active ingredients applied at full recommended dosages.

The density distribution of EC50 values followed a log-normal distribution for all AIs (Figure 3) as commonly observed for many natural phenomena ^30^. A similar distribution of fungicide resistance was observed in field populations of the pathogens *Fusarium graminearum*, *Pyricularia graminis-tritici*, and *Botrytis cinerea* ^49, 54, 16^. For these other fungi, large numbers of isolates were sampled from many different fields distributed across large geographical areas and often over several years. It was not reported in the other studies how intense was the fungicide selection pressure associated with each field collection. We consider it noteworthy that we found a similar distribution of fungicide sensitivities within a single field and during a single growing season in an environment that we expected would select strongly for fungicide resistance. This illustrates well that *Z. tritici* field populations can maintain a high evolutionary potential even after several cycles of strong selection.

The EC50 values showed a heritability of 0.53, consistent with an underlying genetic basis for this trait (Supplemental Table 2). We found a high correlation between the three replications of our experiment, indicating that the data were highly reproducible. Positive and high correlations between EC50 values for propiconazole, epoxiconazole, and metconazole provided evidence for cross-resistance among these DMIs (Table 2). Similar cross-resistance was already reported between propiconazole and different DMIs in *Z. tritici* ^50^ and *Alternaria alternata* ^21^.

In a previous study, Karisto ^25^ estimated, based on the recorded daily temperatures and daily records of precipitation, that two cycles of asexual reproduction occurred during the 43 days separating the C1 and C3 sampling dates. Fungicides were applied twice during this period, at 5 days and 16 days after C1. Despite the evidence for a high heritability and the possibility for two cycles of selection, the percentage of resistant isolates for all individual AIs except prothioconazole was higher in C1 than in C3 (Figure 4). In addition, frequencies of MDR and DDR were higher in C1. This suggests that the fungicide applications occurring after C1 did not create a more resistant fungal population. This counter-intuitive finding was unexpected, so we describe next two different processes that could function alone or together to explain the pattern. Process 1: Escape from selection. In this process, a significant fraction of the *Z. tritici* population was not exposed to a sufficiently high dose of an AI to kill the sensitive strains. The effective period of control for any fungicide ranges between 12-19 days, depending on the AI and temperature ^15^. *Z. tritici* is a necrotroph that continues to colonize the killed leaf tissue as well as naturally senescent leaf tissue and can persist on the lower senesced wheat leaves for many months. We hypothesize that fungicides applied to the top 3-4 green leaf layers of the canopy did not reach lethal concentrations in the senesced lower leaf layers. In this case, ascospores and pycnidiospores produced by fungicide-sensitive isolates in the lower leaf layers would be able to initiate new infections on the upper leaf layers after the fungicide concentrations fell below lethal levels in the treated upper leaves. Process 2: High gene flow from untreated populations. In this process, a large number of airborne ascospores coming from nearby wheat fields that were not treated with fungicides introduced a high frequency of fungicide-sensitive strains into the upper leaf layers of the experimental plots between C1 and C3, diluting the selected fungicide resistant strains. Previous studies showed that airborne ascospores coming from both outside a field and from lower leaves within a field can provide a significant source of inoculum on upper leaf layers during the later stages of an STB epidemic^17,57^.

### Defining a resistance threshold

Using different methods to assign individuals into resistant and sensitive categories yielded different outcomes, though all were highly correlated (R^2^=0.86, *p*-value = 0.006). An approach based on the average EC50 value of each AI dataset was used as a reference point for comparison with other methods. The Scott-Knott and *K*-means (*K*=5) approaches both followed the principle of dividing the dataset into five different categories of fungicide resistance (i.e. HR, R, M, S, and HS). The Scott-Knott method requires the investigators to make an eyeball assessment of where to place the line dividing different categories of resistance, introducing an investigator-specific bias, while the *K*-means (*K*=5) approach separated the isolates based on Euclidian distances. Additional advantages of the *K*-means approach include that it used the fitted values of EC50 and it separated the resistance categories based on a pre-defined logic. The number of isolates defined as resistant using *K*-means (*K*=2) and *K*-means (*K*=5) differed significantly only for bixafen and metconazole, with the 5-category definition always identifying fewer resistant isolates than the 2-category definition. We chose the more conservative *K*-means (*K*=5) definition to identify resistant isolates.

We found that MDR strains are common, occurring in ∼19% of all isolates sampled from this single field. The percentage of MDR dropped from 27% in C1 to 12% in C3 (Figure 4). The most common MDR was to EM. Because we expected that all the isolates were exposed to at least two AIs, we expected to find a relatively high proportion of strains with multidrug resistance, but we also expected that the frequency of MDR strains would increase over time due to additional cycles of selection by fungicides applied as mixtures. Assuming independence between traits, we used the observed frequencies of isolates resistant to individual AIs to calculate the expected frequencies of multidrug resistance phenotypes, and then used a binomial test to determine if the differences were significant. We found that the observed values of most MDRs were significantly higher than expected, suggesting that selection favored the emergence of these MDR phenotypes in this experiment (Figure 7). Interestingly, resistance to BP was not observed very often (8/984) amongst isolates, although this is one of the fungicide combinations that was applied in the field. Applying a mixture of fungicides is expected to postpone the development of resistance to a single AI (Van den Bosch et al., 2014), but may facilitate the emergence of MDR ^2^ by providing a fitness advantage to MDR strains through eliminating competing strains that carry resistance to only one AI.

### The host-fungicide-pathogen (H-F-P) interaction

Wheat cultivars with different genetic backgrounds and expressing different degrees of STB resistance are known to directly impact the establishment of infection and the frequency of sexual and asexual reproduction ^8^. We found abundant evidence that different cultivars hosted populations of isolates expressing different degrees of fungicide sensitivity, i.e. there were significant H-F-P interactions (Figure 5, Supplemental Figure 4, Supplemental Tables 3 and 4). Almost all cultivar-isolate subgroups behaved differentially for AI resistance as indicated by EC50 values, suggesting that the cultivar genotype interacts with the fungicide environment to select the most adapted pathogen isolates. As examples, the isolates sampled from Zinal exhibited a significantly higher EC50 for prothioconazole, while isolates sampled from Runal and Cassiopeia were more resistant to bixafen (Figure 5). A similar pattern of cultivar-specific fungicide resistance emerged when using the *K*-means (*K*=5) definition of resistance to calculate the percentage of resistant isolates on each host cultivar (Figure 6). In this case, Cassiopeia yielded the highest frequency of bixafen-resistant isolates, Zinal hosted the highest frequency of prothioconazole-resistant isolates and Arina hosted the highest frequency of spiroxamine-resistant isolates.

In general, positive relationships were found between the degree of resistance to STB measured on each cultivar and the fungicide resistance of the *Z. tritici* isolates obtained from that same cultivar, though the correlations were significant only for metconazole and bixafen (Table 5). A highly significant correlation (*p*-value < 0.001) was also found between STB resistance and multidrug resistance (Table 5). It is important to note here that the leaves used to obtain the measures of STB resistance are the same leaves that were used to obtain pathogen isolates, so the same pathogen populations were used to make the measurements of host STB resistance and pathogen fungicide resistance. Our findings suggest that more resistant hosts, on average, select for pathogen isolates that are more resistant to fungicides, a pattern that was first described in populations of *Z. tritici* isolated from a resistant wheat cultivar and a susceptible wheat cultivar growing in the same Oregon field that was not treated with fungicides ^59^. Other analyses that included a global collection of 145 *Z. tritici* isolates identified fitness trade-offs between cultivar STB resistance and fungicide sensitivity, with individual isolates that carried higher levels of resistance to particular fungicides being less virulent on particular cultivars ^11^. The new dataset reported here suggests that different hosts selectively favor pathogen isolates carrying resistance to different fungicides. Different cultivars showed significant differences in the percentage of isolates that were resistant to different AIs. For example, Figure 6, which shows the cultivars ordered from most resistant to most susceptible to STB, suggests that Arina favors *Z. tritici* isolates that are more resistant to spiroxamine, Zinal favors isolates that are more resistant to prothioconazole, and Haldor favors isolates that are more resistant to epoxiconazole. This type of information may eventually prove useful for determining which fungicides should be applied to which cultivars to reduce the risk of emergence of fungicide resistance.

The overall MDR percentage did not differ across cultivars, averaging about 19% (Figure 8). The resistant cultivar Cassiopeia hosted more isolates with resistance to the SP and BP combinations, with the BP resistance at a significantly higher frequency compared to other cultivars (Figure 6). This illustrates that specific DDR resistance may also be affected by specific host cultivars.

### Conclusions

Our experiment showed that a high variability for fungicide sensitivity can be maintained in a pathogen population at the field scale even after three treatments with fungicide mixtures. Applications of fungicide mixtures created an environment conducive to the emergence of strains with MDR. More resistant wheat cultivars on average selected for pathogen strains with higher fungicide resistance. There was strong evidence for host-fungicide-pathogen interactions in which particular host genotypes selected for pathogen strains that were more resistant to particular active ingredients, suggesting that some host cultivars may accelerate the emergence of resistance to some fungicides.

