## Supplemental_Table_1 for "Cultivar-specific fungicide resistance emerges during a growing season in field populations of *Zymoseptoria tritici*"

Supplemental Table 1: Concentrations of active ingredients used in a second round of data collection for resistant fungal isolates that needed higher concentrations to calculate their EC50 values.

| Active ingredient | conc. 1 (mg.l-1) | conc. 2 (mg.l-1) | conc. 3 (mg.l-1) | conc. 4 (mg.l-1) | conc. 5 (mg.l-1) | conc. 6 (mg.l-1) | conc. 7 (mg.l-1) | conc. 8 (mg.l-1) |
|---|---|---|---|---|---|---|---|---|
| Bixafen | 0 | 0.001 | 0.05 | 0.3 | 0.5 | 1.5 | 10 | 80 |
| Epoxiconazole | 0 | 5 | 7 | 20 | 50 | 100 | 200 | 250 |
| Metconazole | 0 | 1 | 7 | 20 | 50 | 150 | 200 | 250 |
| Propiconazole | 0 | 1 | 7 | 20 | 50 | 150 | 200 | 300 |
| Prothioconazole | 0 | 1 | 7 | 20 | 50 | 150 | 200 | 300 |
| Spiroxamine | 0 | 0.02 | 0.5 | 0.7 | 2 | 5 | 15 | 100 |
