## Supplemental_Table_2 for "Cultivar-specific fungicide resistance emerges during a growing season in field populations of *Zymoseptoria tritici*"

Supplemental Table 2. Variance component analysis for effects of fungal isolate, host cultivar, and active ingredient on EC50 values. The broad-sense heritability (H2) was calculated based on the genetic vs. Phenotypic

| **Variance Component** | **Estimate** | **Std.Dev** | $H^2$ |
|---|---|---|---|
| Isolate | 0.50 | 0.71 | 0.53 |
| Host cultivar | 0.00 | 0.06 | |
| Fungicide | 2.62 | 1.62 | |
| Replicate | 0.00 | 0.02 | |
| Residual | 0.45 | 0.67 | |
