## Supplemental_Table_3 for "Cultivar-specific fungicide resistance emerges during a growing season in field populations of *Zymoseptoria tritici*"

Supplemental Table 3. Results of modelling the effect of collection time, replicate and host on EC50.

| Active ingredient | source of variation | Df | Sum Sq | Mean Sq | F value | Pr(>F) | |
|---|---|---|---|---|---|---|---|
| Bixafen | **Collection** | **1** | **248.46** | **248.459** | **230.89** | **<2e-16** | *** |
| | Replications | 1 | 0.01 | 0.009 | 0.01 | 0.9261 | |
| | **Host** | **14** | **151.65** | **10.832** | **10.07** | **<2e-16** | *** |
| | Residuals | 2929 | 3151.81 | 1.076 | | | |
| Epoxiconazole | **Collection** | **1** | **275.4** | **275.358** | **197.21** | **<2e-16** | *** |
| | Replications | 1 | 0.4 | 0.374 | 0.27 | 0.605 | |
| | **Host** | **14** | **95.5** | **6.821** | **4.88** | **4.99E-09** | *** |
| | Residuals | 2828 | 3948.7 | 1.396 | | | |
| Metconazole | **Collection** | **1** | **72.02** | **36.01** | **31.99** | **1.85E-14** | *** |
| | **Replications** | **1** | **4.43** | **4.428** | **3.93** | **4.74E-02** | * |
| | **Host** | **14** | **97.88** | **6.991** | **6.21** | **2.35E-12** | *** |
| | Residuals | 2741 | 3085.33 | 1.126 | | | |
| Propiconazole | **Collection** | **1** | **23.99** | **23.994** | **32.77** | **1.15E-08** | *** |
| | Replications | 1 | 0.88 | 0.8777 | 1.20 | 0.2737 | |
| | **Host** | **14** | **37.48** | **2.6774** | **3.66** | **4.41E-06** | *** |
| | **Residuals** | **2892** | **2117.77** | **0.7323** | | | |
| Prothioconazole | **Collection** | **1** | **0.95** | **0.95282** | **8.16** | **0.004308** | ** |
| | Replications | 1 | 0 | 0.0013 | 0.01 | 0.915909 | |
| | **Host** | **14** | **20.67** | **1.47674** | **12.65** | **<2e-16** | *** |
| | Residuals | 2865 | 334.44 | 0.11673 | | | |
| Spiroxamine | Collection | 1 | 0.23 | 0.23289 | 0.79 | 0.373893 | |
| | **Replications** | **1** | **2.9** | **2.90118** | **9.85** | **0.001713** | ** |
| | **Host** | **14** | **26.67** | **1.90515** | **6.47** | **4.98E-13** | *** |
| | Residuals | 2897 | 853.04 | 0.29446 | | | |
