## Supplemental_Table_4 for "Cultivar-specific fungicide resistance emerges during a growing season in field populations of *Zymoseptoria tritici*"

Supplemental Table 4. Results of modelling the effect of collection time and host on mean EC50 across replicates for each AI separately.

Bixafen

| | Df | Sum Sq | Mean Sq | F value | Pr(>F) | |
|---|---|---|---|---|---|---|
| **collection** | **1** | **88.15** | **88.152** | **77.77** | **< 2.20E-16** | *** |
| **Host** | **14** | **60.68** | **4.334** | **3.82** | **2.39E-06** | *** |
| **collection:Host** | **14** | **51.52** | **3.68** | **3.25** | **4.59E-05** | *** |
| Residuals | 960 | 1088.11 | 1.133 | | | |

Epoxiconazole

| | Df | Sum Sq | Mean Sq | F value | Pr(>F) | |
|---|---|---|---|---|---|---|
| **collection** | **1** | **109.07** | **109.072** | **78.54** | **< 2.20E-16** | *** |
| Host | 14 | 32.48 | 2.32 | 1.67 | 0.05629 | . |
| collection:Host | 14 | 30.82 | 2.201 | 1.59 | 0.07713 | . |
| Residuals | 947 | 1315.15 | 1.389 | | | |

Metconazole

| | Df | SumSq | MeanSq | F | p | |
|---|---|---|---|---|---|---|
| **collection** | **1** | **23.26** | **23.2583** | **18.15** | **2.25E-05** | *** |
| **Host** | **14** | **37.84** | **2.7028** | **2.11** | **0.0096611** | ** |
| **collection:Host** | **14** | **50.93** | **3.638** | **2.84** | **0.0003447** | *** |
| Residuals | 910 | 1166.06 | 1.2814 | | | |

Propiconazole

| | Df | SumSq | MeanSq | F | p | |
|---|---|---|---|---|---|---|
| **collection** | **1** | **7.9** | **7.903** | **6.38** | **0.01172** | * |
| Host | 14 | 21.67 | 1.5477 | 1.25 | 0.23367 | |
| collection:Host | 14 | 20.36 | 1.4544 | 1.17 | 0.28989 | |
| Residuals | 951 | 1178.43 | 1.239 | | | |

Prothioconazole

| | Df | SumSq | MeanSq | F | p | |
|---|---|---|---|---|---|---|
| collection | 1 | 0.79 | 0.78804 | 1.29 | 0.25715 | |
| **Host** | **14** | **17.55** | **1.25336** | **2.04** | **0.01265** | * |
| collection:Host | 14 | 13.06 | 0.93313 | 1.52 | 0.09638 | . |
| Residuals | 960 | 588.47 | 0.61299 | | | |

Spiroxamine

| | Df | SumSq | MeanSq | F | p | |
|---|---|---|---|---|---|---|
| collection | 1 | 0.51 | 0.5086 | 0.84 | 0.35888 | |
| **Host** | **14** | **19.37** | **1.3836** | **2.29** | **4.33E-03** | ** |
| **collection:Host** | **14** | **29.77** | **2.1261** | **3.52** | **1.13E-05** | *** |
| Residuals | 959 | 578.85 | 0.6036 | | | |
