## Supplemental_Table_5 for "Cultivar-specific fungicide resistance emerges during a growing season in field populations of *Zymoseptoria tritici*"

Supplemental Table 5. Result of modelling the effect of collection time and host on the frequency of resistance to each AI, and the frequency of MDR and DDR.

| Bixafen | | Df | Deviance | Residual Df | Residual Deviance | p | |
|---|---|---|---|---|---|---|---|
| | NULL | **989** | **458.16** | | | | |
| | **collection** | **1** | **23.909** | **988** | **434.25** | **1.01E-06** | *** |
| | Host | 14 | 16.97 | 974 | 417.28 | 0.2578 | |

| Epoxiconazole | | Df | Deviance | Residual Df | Residual Deviance | p | |
|---|---|---|---|---|---|---|---|
| | NULL | 976 | 810.02 | | | | |
| | **collection** | **1** | **58.857** | **975** | **751.16** | **1.70E-14** | *** |
| | Host | 14 | 16.876 | 961 | 734.29 | 0.262861 | |

Metconazole

| | | Df | Deviance | Residual Df | Residual Deviance | p | |
|---|---|---|---|---|---|---|---|
| | NULL | 939 | 780.68 | | | | |
| | **collection** | **1** | **21.939** | **938** | **758.74** | **2.81E-06** | *** |
| | Host | 14 | 16.704 | 924 | 742.04 | 0.2723 | |

| Propiconazole | | Df | Deviance | Residual Df | Residual Deviance | p | |
|---|---|---|---|---|---|---|---|
| | NULL | 980 | 740.68 | | | | |
| | **collection** | **1** | **14.876** | **979** | **725.8** | **0.0001148** | *** |
| | Host | 14 | 10.344 | 965 | 715.46 | 0.7366068 | |

| Prothioconazole | | Df | Deviance | Residual Df | Residual Deviance | p | |
|---|---|---|---|---|---|---|---|
| | NULL | 989 | 526.18 | | | | |
| | collection | 1 | 0.569 | 988 | 525.62 | 0.4506 | |
| | **Host** | **14** | **60.32** | **974** | **465.29** | **1.03E-07** | *** |

| Spiroxamine | | Df | Deviance | Residual Df | Residual Deviance | p | |
|---|---|---|---|---|---|---|---|
| | NULL | 988 | 923.19 | | | | |
| | **collection** | **1** | **5.394** | **987** | **917.8** | **0.0202093** | * |
| | **Host** | **14** | **37.316** | **973** | **8.80E+02** | **0.0006607** | *** |

| MDR (≥2 ai) | | Df | Deviance | Residual Df | Residual Deviance | p | |
|---|---|---|---|---|---|---|---|
| | NULL | 997 | 951.01 | | | | |
| | **collection** | **1** | **33.272** | **996** | **917.74** | **8.01E-09** | *** |
| | Host | 14 | 9.859 | 982 | 907.88 | 0.7724 | |

| DDR (all 2 AI combinations) | | Df | Deviance | Residual Df | Residual Deviance | p |
|---|---|---|---|---|---|---|
| | NULL | 997 | 517.26 | | | |
| | collection | 1 | 0.9125 | 996 | 516.35 | 0.3394 |
| | Host.resOrder | 14 | 20.1586 | 982 | 496.19 | 0.1252 |

| EM (Epoxiconazole+Metconazole) | | Df | Deviance | Residual Df | Residual Deviance | p | |
|---|---|---|---|---|---|---|---|
| | NULL | 922 | 558.16 | | | | |
| | **collection** | **1** | **45.213** | **921** | **512.95** | **1.77E-11** | *** |
| | Host | 14 | 13.778 | 907 | 499.17 | 0.4664 | |

| BP (Bixafen+Prothioconazole) | | Df | Deviance | Residual Df | Residual Deviance | p | |
|---|---|---|---|---|---|---|---|
| | NULL | 983 | 92.93 | | | | |
| | **collection** | **1** | **3.9236** | **982** | **89.006** | **0.04761** | * |
| | **Host** | **14** | **26.1688** | **968** | **62.837** | **0.02464** | * |

| SP (Spiroxamine+Prothioconazole) | | Df | Deviance | Residual Df | Residual Deviance | p |
|---|---|---|---|---|---|---|
| | NULL | 982 | 171.66 | | | |
| | collection | 1 | 1.1266 | 981 | 170.53 | 0.2885 |
| | Host.resOrder | 14 | 20.3863 | 967 | 150.14 | 0.1184 |
