## Supplemental_Table_6 for "Cultivar-specific fungicide resistance emerges during a growing season in field populations of *Zymoseptoria tritici*"

Supplemental Table 6. Analysis of variance of normlaized and log-transfomed EC50 by collection,host, AI and hostxAI interaction (Host:ai)

| | Df | SumSq | MeanSq | F | p |
|---|---|---|---|---|---|
| collection | 1 | 125.5 | 125.52 | 116.64 | <2.00E-16 |
| Host | 14 | 67.5 | 4.82 | 4.48 | 4.35E-08 |
| AI | 5 | 462.1 | 92.41 | 85.87 | <2.00E-16 |
| Host:AI | 70 | 125.3 | 1.79 | 1.66 | 0.000461 |
| Residuals | 5775 | 6214.8 | 1.08 | | |
