## Supplemental_Table_7 for "Cultivar-specific fungicide resistance emerges during a growing season in field populations of *Zymoseptoria tritici*"

Supplemental Table 7: Correlation between EC50 cultivar based (ie., mean EC50 value of isolates hosted by a certain cultivar), Z. tritici pathogenicity (PLACAL, pycnidia density, and ρ-leaf) and the percentage of Dual- and Multi- drug resistant isolates per cultivars. Correlation coefficients are in the upper diagonal and p-values are in the lower diagonal. P-values are only provided for strictly independent variables (e.g. Ec_Bix, Res%_Bix and Res%_BP are not independent variables).

| | PLACL | ρlesions | ρleaf | EC_Bix | EC_Epo | EC_Met | EC_Propi | EC_Prth | EC_Spr | % Res_Bix | % Res_Epo | % Res_Met | % Res_Propi | % Res_Prth | % Res_Spr | % Res_MDR | % Res_SP | % Res_BP | % Res_EM |
|---|---|---|---|---|---|---|---|---|---|---|---|---|---|---|---|---|---|---|---|
| PLACL | | 0.25 | 0.68 | -0.27 | -0.06 | -0.61 | 0.22 | -0.17 | -0.01 | -0.24 | 0.03 | -0.43 | 0.21 | -0.14 | -0.25 | -0.16 | -0.48 | -0.34 | -0.30 |
| ρlesions | 0.37 | | 0.71 | -0.41 | -0.31 | -0.33 | 0.19 | 0.21 | -0.56 | -0.12 | -0.30 | -0.25 | 0.24 | 0.12 | -0.65 | **-0.82** | 0.14 | 0.25 | -0.12 |
| ρleaf | 0.01 | 0.00 | | -0.44 | -0.26 | -0.59 | 0.15 | -0.07 | -0.36 | -0.22 | -0.17 | -0.40 | 0.22 | -0.11 | -0.48 | **-0.60** | -0.18 | -0.05 | -0.22 |
| EC_Bix | 0.34 | 0.12 | 0.10 | | 0.45 | 0.77 | 0.47 | 0.13 | 0.21 | 0.88 | 0.37 | 0.77 | 0.40 | 0.18 | 0.23 | 0.68 | 0.51 | 0.62 | 0.56 |
| EC_Epo | 0.84 | 0.26 | 0.36 | 0.09 | | 0.35 | 0.42 | -0.34 | 0.43 | 0.26 | 0.86 | 0.31 | 0.41 | -0.36 | 0.45 | 0.65 | -0.05 | 0.03 | 0.68 |
| EC_Met | 0.02 | 0.24 | 0.02 | 0.001 | 0.20 | | 0.23 | 0.04 | 0.05 | 0.67 | 0.25 | 0.89 | 0.12 | 0.03 | 0.23 | 0.50 | 0.59 | 0.63 | 0.55 |
| EC_Propi | 0.43 | 0.50 | 0.59 | 0.08 | 0.12 | 0.40 | | 0.00 | 0.30 | 0.63 | 0.23 | 0.37 | 0.88 | -0.05 | -0.04 | 0.25 | 0.24 | 0.40 | 0.54 |
| EC_Prth | 0.55 | 0.45 | 0.79 | 0.64 | 0.22 | 0.88 | 1.00 | | -0.20 | 0.13 | -0.43 | 0.03 | -0.05 | 0.94 | -0.30 | -0.22 | 0.41 | 0.29 | -0.22 |
| EC_Spr | 0.99 | 0.03 | 0.19 | 0.45 | 0.11 | 0.87 | 0.28 | 0.47 | | 0.09 | 0.16 | 0.06 | 0.14 | -0.16 | 0.86 | 0.48 | 0.03 | -0.08 | 0.33 |
| % Res_Bix | 0.40 | 0.67 | 0.44 | NA | 0.35 | 0.01 | 0.01 | 0.64 | 0.75 | | 0.21 | 0.76 | 0.60 | 0.17 | 0.06 | 0.45 | 0.62 | 0.76 | 0.59 |
| % Res_Epo | 0.91 | 0.28 | 0.55 | 0.17 | NA | 0.36 | 0.41 | 0.11 | 0.56 | 0.45 | | 0.22 | 0.33 | -0.43 | 0.26 | 0.65 | -0.22 | -0.11 | 0.59 |
| % Res_Met | 0.11 | 0.37 | 0.13 | 0.00 | 0.26 | NA | 0.17 | 0.91 | 0.84 | 0.00 | 0.43 | | 0.22 | -0.06 | 0.14 | 0.50 | 0.52 | 0.62 | 0.71 |
| % Res_Propi | 0.45 | 0.40 | 0.44 | 0.14 | 0.13 | 0.68 | NA | 0.86 | 0.62 | 0.02 | 0.23 | 0.43 | | -0.08 | -0.13 | 0.24 | 0.06 | 0.28 | 0.50 |
| % Res_Prth | 0.62 | 0.68 | 0.70 | 0.53 | 0.18 | 0.92 | 0.85 | NA | 0.58 | 0.54 | 0.11 | 0.84 | 0.78 | | -0.19 | -0.20 | 0.48 | 0.34 | -0.36 |
| % Res_Spr | 0.37 | 0.01 | 0.07 | 0.41 | 0.09 | 0.41 | 0.88 | 0.28 | NA | 0.83 | 0.36 | 0.62 | 0.64 | 0.49 | | 0.47 | 0.16 | 0.02 | 0.31 |
| % Res_MDR | 0.58 | **0.000** | **0.02** | 0.01 | 0.01 | 0.06 | 0.36 | 0.43 | 0.07 | NA | NA | NA | NA | NA | NA | | -0.11 | -0.08 | 0.52 |
| % Res_SP | 0.07 | 0.62 | 0.52 | 0.05 | 0.86 | 0.02 | 0.38 | NA | NA | 0.01 | 0.44 | 0.05 | 0.83 | NA | NA | NA | | 0.93 | 0.24 |
| % Res_BP | 0.22 | 0.37 | 0.85 | NA | 0.91 | 0.01 | 0.14 | NA | 0.78 | NA | 0.69 | 0.01 | 0.31 | NA | 0.93 | NA | NA | | 0.36 |
| % Res_EM | 0.28 | 0.67 | 0.42 | 0.03 | NA | NA | 0.04 | 0.44 | 0.24 | 0.02 | NA | NA | 0.06 | 0.19 | 0.26 | NA | 0.38 | 0.19 | |
