## Supplemental_Table_8 for "Cultivar-specific fungicide resistance emerges during a growing season in field populations of *Zymoseptoria tritici*"

Supplemental Table 8: Number of isolates that show multi-drug resistance and binomial test to investigate if these are different to expectation. B=bixafen, E=epoxiconazole, M=metconazole, P=prothioconazole, R=propiconazole, and S=spiroxamine.

| order | n.ai | combo | n.mdr | tot.mdr | exp.prop | prop.mdr | prop.diff | pval | adj.pval.bonf | |
|---|---|---|---|---|---|---|---|---|---|---|
| 1 | 2 | E,M | 83 | 923 | 0.0200 | 0.0900 | 0.0700 | 1.64E-27 | 2.47E-26 | *** |
| 2 | 2 | E,R | 72 | 968 | 0.0200 | 0.0700 | 0.0600 | 5.50E-23 | 8.25E-22 | *** |
| 3 | 2 | E,B | 34 | 971 | 0.0100 | 0.0400 | 0.0300 | 4.69E-11 | 7.03E-10 | *** |
| 4 | 2 | E,P | 19 | 971 | 0.0100 | 0.0200 | 0.0100 | 0.01827826 | 0.27417382 | ns |
| 5 | 2 | E,S | 73 | 970 | 0.0300 | 0.0800 | 0.0500 | 1.38E-15 | 2.07E-14 | *** |
| 6 | 2 | M,R | 80 | 930 | 0.0200 | 0.0900 | 0.0700 | 2.17E-29 | 3.26E-28 | *** |
| 7 | 2 | M,B | 39 | 934 | 0.0100 | 0.0400 | 0.0300 | 9.00E-15 | 1.35E-13 | *** |
| 8 | 2 | M,P | 22 | 932 | 0.0100 | 0.0200 | 0.0100 | 0.00120222 | 0.01803334 | * |
| 9 | 2 | M,S | 76 | 932 | 0.0300 | 0.0800 | 0.0600 | 5.51E-18 | 8.26E-17 | *** |
| 10 | 2 | R,B | 36 | 974 | 0.0100 | 0.0400 | 0.0300 | 4.31E-14 | 6.46E-13 | *** |
| 11 | 2 | R,P | 14 | 974 | 0.0100 | 0.0100 | 0.0100 | 0.1295613 | 1 | ns |
| 12 | 2 | R,S | 71 | 974 | 0.0200 | 0.0700 | 0.0500 | 1.10E-17 | 1.64E-16 | *** |
| 13 | 2 | B,P | 8 | 984 | 0.0000 | 0.0100 | 0.0000 | 0.09941216 | 1 | ns |
| 14 | 2 | B,S | 44 | 982 | 0.0100 | 0.0400 | 0.0300 | 1.26E-14 | 1.90E-13 | *** |
| 15 | 2 | P,S | 17 | 983 | 0.0100 | 0.0200 | 0.0000 | 0.26134812 | 1 | ns |
| 16 | 3 | E,M,R | 56 | 918 | 0.0027 | 0.0610 | 0.0583 | 1.27E-55 | 2.53E-54 | *** |
| 17 | 3 | E,M,B | 25 | 918 | 0.0013 | 0.0272 | 0.0259 | 1.39E-24 | 2.78E-23 | *** |
| 18 | 3 | E,M,P | 9 | 917 | 0.0016 | 0.0098 | 0.0082 | 2.10E-05 | 0.00042006 | *** |
| 19 | 3 | E,M,S | 54 | 917 | 0.0037 | 0.0589 | 0.0551 | 3.30E-45 | 6.61E-44 | *** |
| 20 | 3 | E,R,B | 24 | 962 | 0.0011 | 0.0249 | 0.0238 | 2.80E-24 | 5.59E-23 | *** |
| 21 | 3 | E,R,P | 9 | 962 | 0.0014 | 0.0094 | 0.0080 | 9.46E-06 | 0.00018926 | *** |
| 22 | 3 | E,R,S | 45 | 962 | 0.0032 | 0.0468 | 0.0436 | 2.16E-36 | 4.32E-35 | *** |
| 23 | 3 | E,B,P | 5 | 966 | 0.0007 | 0.0052 | 0.0045 | 0.00054776 | 0.0109553 | * |
| 24 | 3 | E,B,S | 24 | 964 | 0.0016 | 0.0249 | 0.0233 | 7.54E-21 | 1.51E-19 | *** |
| 25 | 3 | E,P,S | 8 | 964 | 0.0019 | 0.0083 | 0.0064 | 0.00066525 | 0.01330501 | * |
| 26 | 3 | M,R,B | 31 | 924 | 0.0011 | 0.0335 | 0.0324 | 9.42E-35 | 1.88E-33 | *** |
| 27 | 3 | M,R,P | 10 | 923 | 0.0014 | 0.0108 | 0.0095 | 8.61E-07 | 1.72E-05 | *** |
| 28 | 3 | M,R,S | 55 | 924 | 0.0032 | 0.0595 | 0.0563 | 1.34E-49 | 2.69E-48 | *** |
| 29 | 3 | M,B,P | 6 | 928 | 0.0007 | 0.0065 | 0.0058 | 4.71E-05 | 0.00094215 | *** |
| 30 | 3 | M,B,S | 32 | 927 | 0.0016 | 0.0345 | 0.0329 | 1.34E-31 | 2.69E-30 | *** |
| 31 | 3 | M,P,S | 10 | 926 | 0.0019 | 0.0108 | 0.0089 | 1.75E-05 | 0.00034979 | *** |
| 32 | 3 | R,B,P | 5 | 969 | 0.0006 | 0.0052 | 0.0046 | 0.00028539 | 0.00570788 | ** |
| 33 | 3 | R,B,S | 31 | 968 | 0.0014 | 0.0320 | 0.0307 | 1.28E-31 | 2.55E-30 | *** |
| 34 | 3 | R,P,S | 9 | 968 | 0.0017 | 0.0093 | 0.0076 | 4.54E-05 | 0.00090736 | *** |
| 35 | 3 | B,P,S | 5 | 977 | 0.0008 | 0.0051 | 0.0043 | 0.00137262 | 0.02745238 | * |
| 36 | 4 | E,M,R,B | 21 | 913 | 0.00016365 | 0.0230011 | 0.022837 | 6.20E-38 | 9.31E-37 | *** |
| 37 | 4 | E,M,R,P | 7 | 912 | 0.00019853 | 0.00767544 | 0.007477 | 1.06E-09 | 1.59E-08 | *** |
| 38 | 4 | E,M,R,S | 39 | 913 | 0.00046996 | 0.04271632 | 0.042246 | 6.73E-62 | 1.01E-60 | *** |
| 39 | 4 | E,M,B,P | 3 | 913 | 9.76E-05 | 0.00328587 | 0.003188 | 0.00010985 | 0.00164779 | ** |
| 40 | 4 | E,M,B,S | 19 | 912 | 0.00023095 | 0.02083333 | 0.020602 | 7.85E-31 | 1.18E-29 | *** |
| 41 | 4 | E,M,P,S | 8 | 911 | 0.00028017 | 0.00878156 | 0.008501 | 3.46E-10 | 5.19E-09 | *** |
| 42 | 4 | E,R,B,P | 4 | 957 | 8.39E-05 | 0.00417973 | 0.004096 | 1.62E-06 | 2.43E-05 | *** |
| 43 | 4 | E,R,B,S | 19 | 956 | 0.00019869 | 0.01987448 | 0.019676 | 1.13E-31 | 1.70E-30 | *** |
| 44 | 4 | E,R,P,S | 7 | 956 | 0.00024103 | 0.00732218 | 0.007081 | 5.48E-09 | 8.22E-08 | *** |
| 45 | 4 | E,B,P,S | 3 | 959 | 0.00011845 | 0.00312826 | 0.00301 | 0.00022374 | 0.00335611 | ** |
| 46 | 4 | M,R,B,P | 4 | 919 | 8.42E-05 | 0.00435256 | 0.004268 | 1.39E-06 | 2.09E-05 | *** |
| 47 | 4 | M,R,B,S | 27 | 919 | 0.00019924 | 0.02937976 | 0.029181 | 6.51E-49 | 9.76E-48 | *** |
| 48 | 4 | M,R,P,S | 8 | 918 | 0.0002417 | 0.0087146 | 0.008473 | 1.16E-10 | 1.74E-09 | *** |
| 49 | 4 | M,B,P,S | 5 | 922 | 0.00011878 | 0.00542299 | 0.005304 | 1.19E-07 | 1.78E-06 | *** |
| 50 | 4 | R,B,P,S | 4 | 963 | 0.00010218 | 0.00415369 | 0.004052 | 3.59E-06 | 5.38E-05 | *** |
| 51 | 5 | E,M,R,B,P | 3 | 908 | 1.22E-05 | 0.00330397 | 0.003292 | 2.26E-07 | 1.35E-06 | *** |
| 52 | 5 | E,M,R,B,S | 17 | 908 | 2.90E-05 | 0.01872247 | 0.018694 | 3.24E-42 | 1.94E-41 | *** |
| 53 | 5 | E,M,R,P,S | 7 | 907 | 3.51E-05 | 0.00771775 | 0.007683 | 6.29E-15 | 3.77E-14 | *** |
| 54 | 5 | E,M,B,P,S | 3 | 907 | 1.73E-05 | 0.00330761 | 0.00329 | 6.30E-07 | 3.78E-06 | *** |
| 55 | 5 | E,R,B,P,S | 3 | 951 | 1.49E-05 | 0.00315457 | 0.00314 | 4.63E-07 | 2.78E-06 | *** |
| 56 | 5 | M,R,B,P,S | 4 | 914 | 1.49E-05 | 0.00437637 | 0.004361 | 1.41E-09 | 8.43E-09 | *** |
