## Supplementary figures and images for "Cultivar-specific fungicide resistance emerges during a growing season in field populations of *Zymoseptoria tritici*"

### Supplemental_Fig_1

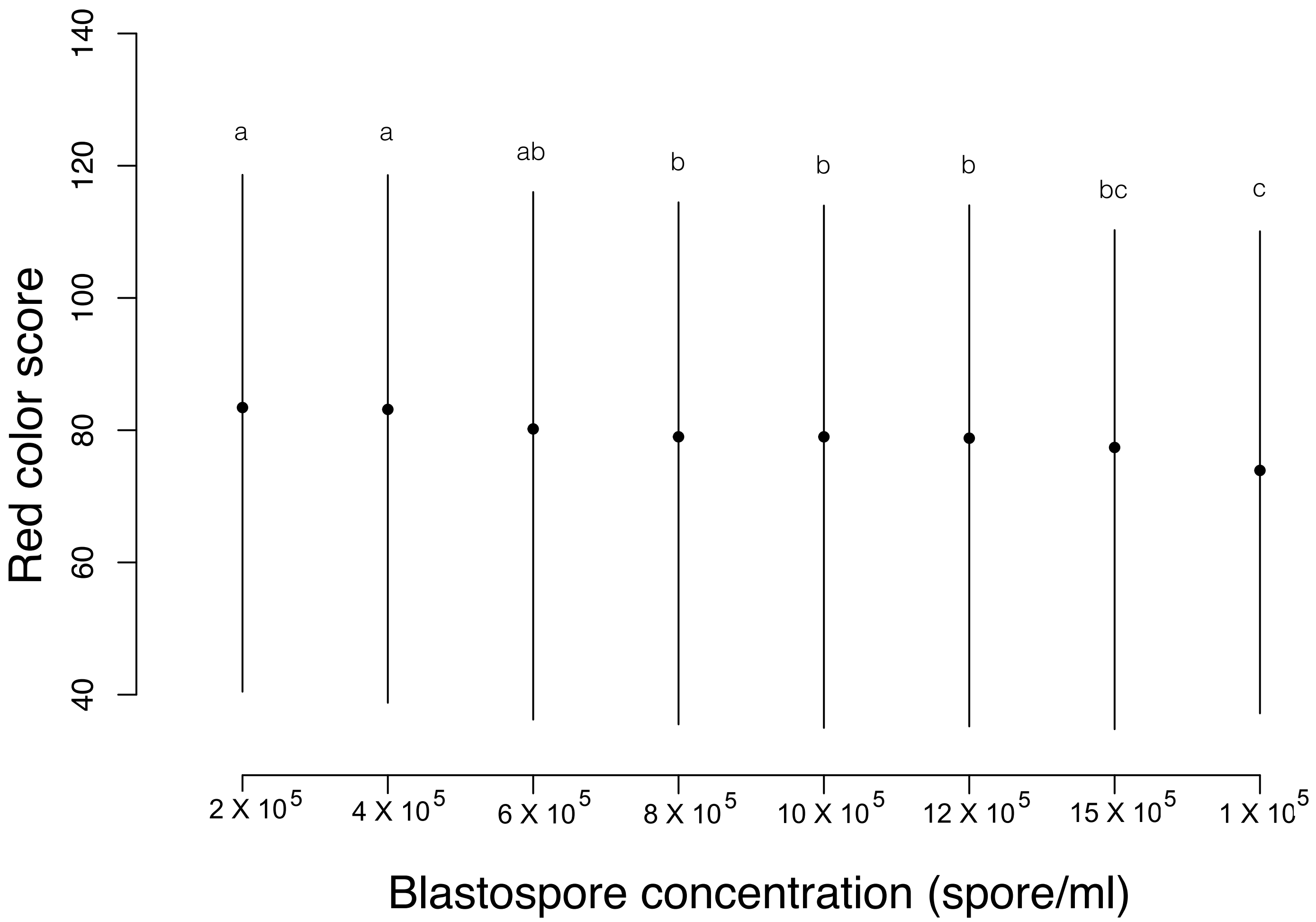

### Supplemental_Fig_2

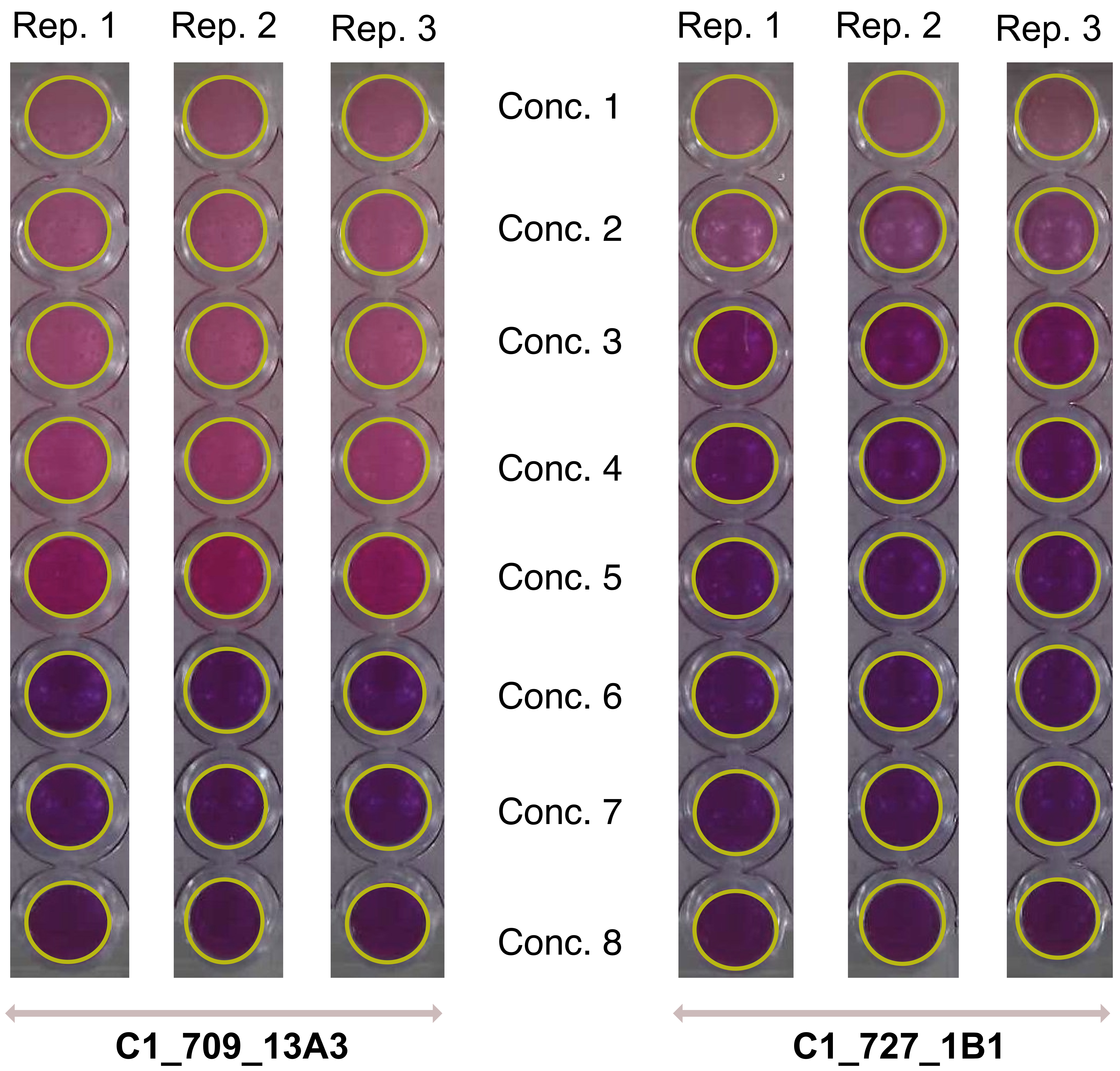

### Supplemental_Fig_3

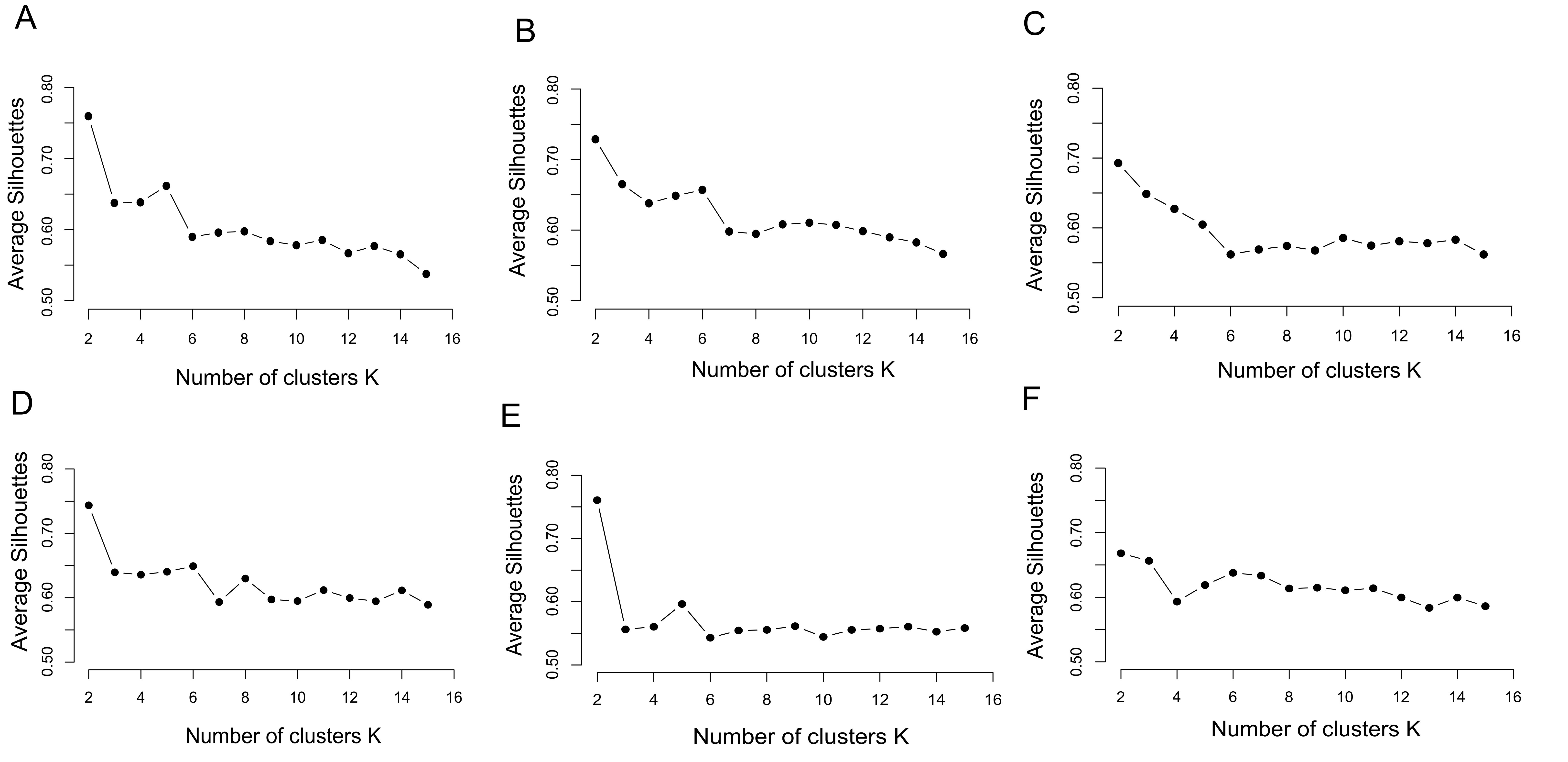

### Supplemental_Fig_4

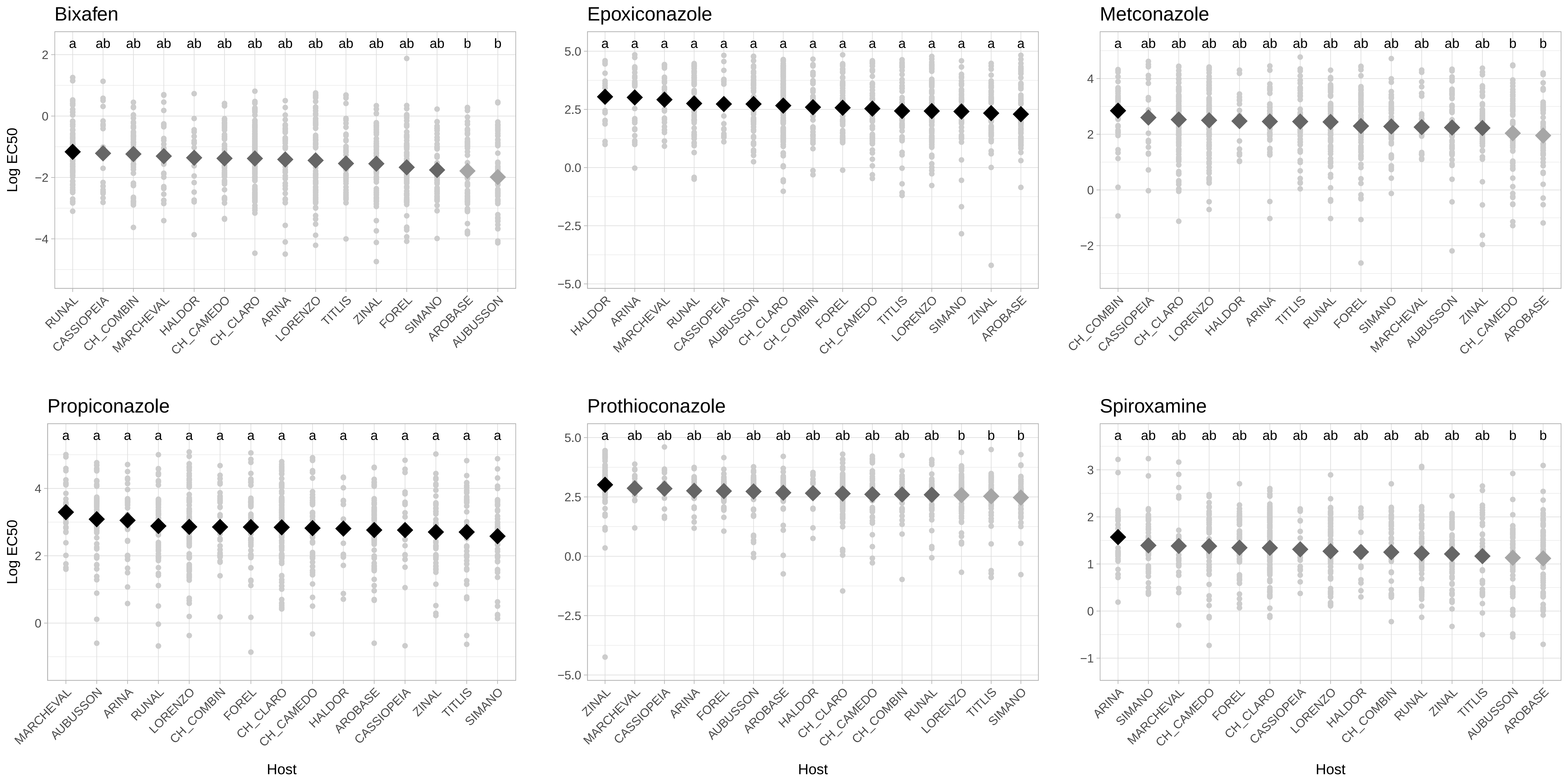
