## Supplemental_File_1 for "Cultivar-specific fungicide resistance emerges during a growing season in field populations of *Zymoseptoria tritici*"

```
## Talas et al. 2024. Cultivar-specific fungicide resistance emerges
during a growing season in ##field populations of Zymoseptoria
tritici##
# this script was modified based on Stewart & McDonald 2014#####
#! /usr/bin/env bash


# Change text colour:
#tput setaf 2 # Change font colour to green.
#tput setab 0 # Change background colour to black
#tput sgr 0 # Change font colour back to normal.
tput bold # Make text bold

echo "$(tput setab 0) $(tput setaf 2)Enter the name of the file
containing your sample names, followed by [ENTER]:$(tput sgr 0)"

read input # Get user input and store it in variable called 'input'

# Add .txt to input file if not already present:
if [[ "${input:(-4)}" != ".txt" ]]
        then
                input=$input".txt"
        fi

# Check if input file is present in working directory:

files=(*)
#echo ${files[@]}
if [[ ${files[*]} =~ $input ]]
        then
        echo 'File found'
else
        directory=$(pwd)

                echo "$(tput setab 0) $(tput setaf 1)File "$input"
not found. Check file is present in your working directory. Your
present working directory is '$directory'. To change youe working
directory type: cd /Path/to/your/directory then hit <ENTER>$(tput
sgr 0)"

                exit
fi



# Make array from input file:

#\015
#\012
#\x0a - hex
#\n
#\r

#fileItemString=$(cat  $input |tr "\n" " ")
```

```
fileItemString=$(cat  $input |tr [:cntrl:] " ")
names=($fileItemString)
length=${#names[@]}

# Ask user to confirm that input file looks OK:

echo "$(tput setab 0) $(tput setaf 2)Your file contains $length
samples and looks like:$(tput sgr 0)"
echo
echo ${names[0]}
echo ${names[1]}
echo "."
echo "."
echo "."
echo ${names[$length-1]}
echo
tput bold

read -p "$(tput setab 0) $(tput setaf 2)Does this look correct? Type
n to exit or y to continue$(tput sgr 0)_" yn # Get user input

case $yn in
        # If user types 'n', show message and exit:
        [Nn]* ) echo "$(tput setab 0) $(tput setaf 1)Fix your file
and try again$(tput sgr 0)"; exit;;
esac

# Get output file name from user:

tput bold

echo "$(tput setab 0) $(tput setaf 2)Enter the name of your output
file, followed by [ENTER]:$(tput sgr 0)"

read output

# Add .tex to output file if not already present:
if [[ "${output:(-4)}" != ".tex" ]]
then
#echo "not .tex"
output=$output".tex"
fi


####perPage=8 # Number of leaves per page####

####pages=$(( ($length + ($perPage -1) ) / $perPage)) # Round up to
nearest whole page.####


### Begin making .tex script ###

## Head Stuff
```

```
echo "\documentclass[a4paper]{minimal}
\usepackage[margin=0cm]{geometry}
\usepackage{auto-pst-pdf,pst-barcode}" > latexTestScript.tex

echo '\\begin{document}' >> latexTestScript.tex # "\" needs to be
quoted. \b = backspace. without quotes "egin{document}" is printed.

## Lines


#pages=6

num=0
for i in $(seq "$pages");
do
 #echo Number: "$i"

#done

echo '\\begin{center}' >> latexTestScript.tex

echo '\\begin{pspicture}[shift=*](20,29)' >> latexTestScript.tex

echo '\psline(0.4,28.26)(19.4,28.26) % top line
\psline(0.4,0.7)(19.4,0.7) % Bottom line
\psline(0.4,0.7)(0.4,28.26) % vertical right line
\psline(19.4,0.7)(19.4,28.26) % vertical left line



% QR Codes' >> latexTestScript.tex

## QR Codes
##column1
if [ $length -gt $num ]
then

texName=$(echo ${names[$num]}  | awk '{gsub( /[#$%&_{}]/, "\\\\&");
print $0}')

echo '\\rput(0.7,0.9){\psbarcode{'${names[$num]}'}{width=0.5
height=0.5}{qrcode}} % Top QR Code
\\rput[Bl](0.7,2.3){'${texName}'}' >> latexTestScript.tex

num=$(($num+1))

fi

if [ $length -gt $num ]
then

texName=$(echo ${names[$num]}  | awk '{gsub( /[#$%&_{}]/, "\\\\&");
```

```
print $0}')

echo '\\rput(0.7,3.02){\psbarcode{'${names[$num]}'}{width=0.5
height=0.5}{qrcode}} % 2nd QR Code
\\rput[Bl](0.7,4.42){'${texName}'}' >> latexTestScript.tex

num=$(($num+1))

fi

if [ $length -gt $num ]
then

texName=$(echo ${names[$num]}  | awk '{gsub( /[#$%&_{}]/, "\\\\&");
print $0}')

echo '\\rput(0.7,5.14){\psbarcode{'${names[$num]}'}{width=0.5
height=0.5}{qrcode}} % 3rd QR Code
\\rput[Bl](0.7,6.54){'${texName}'}' >> latexTestScript.tex

num=$(($num+1))

fi

if [ $length -gt $num ]
then

texName=$(echo ${names[$num]}  | awk '{gsub( /[#$%&_{}]/, "\\\\&");
print $0}')

echo '\\rput(0.7,7.26){\psbarcode{'${names[$num]}'}{width=0.5
height=0.5}{qrcode}} % 4th QR Code
\\rput[Bl](0.7,8.66){'${texName}'}' >> latexTestScript.tex

num=$(($num+1))

fi

if [ $length -gt $num ]
then

texName=$(echo ${names[$num]}  | awk '{gsub( /[#$%&_{}]/, "\\\\&");
print $0}')

echo '\\rput(0.7,9.38){\psbarcode{'${names[$num]}'}{width=0.5
height=0.5}{qrcode}} % 4th QR Code
\\rput[Bl](0.7,10.78){'${texName}'}' >> latexTestScript.tex

num=$(($num+1))

fi

if [ $length -gt $num ]
then
```

```
texName=$(echo ${names[$num]}  | awk '{gsub( /[#$%&_{}]/, "\\\\&");
print $0}')

echo '\\rput(0.7,11.5){\psbarcode{'${names[$num]}'}{width=0.5
height=0.5}{qrcode}} % 4th QR Code
\\rput[Bl](0.7,12.9){'${texName}'}' >> latexTestScript.tex

num=$(($num+1))

fi

if [ $length -gt $num ]
then

texName=$(echo ${names[$num]}  | awk '{gsub( /[#$%&_{}]/, "\\\\&");
print $0}')

echo '\\rput(0.7,13.62){\psbarcode{'${names[$num]}'}{width=0.5
height=0.5}{qrcode}} % 4th QR Code
\\rput[Bl](0.7,15.02){'${texName}'}' >> latexTestScript.tex

num=$(($num+1))

fi

if [ $length -gt $num ]
then

texName=$(echo ${names[$num]}  | awk '{gsub( /[#$%&_{}]/, "\\\\&");
print $0}')

echo '\\rput(0.7,15.74){\psbarcode{'${names[$num]}'}{width=0.5
height=0.5}{qrcode}} % 4th QR Code
\\rput[Bl](0.7,17.14){'${texName}'}' >> latexTestScript.tex

num=$(($num+1))

fi

if [ $length -gt $num ]
then

texName=$(echo ${names[$num]}  | awk '{gsub( /[#$%&_{}]/, "\\\\&");
print $0}')

echo '\\rput(0.7,17.86){\psbarcode{'${names[$num]}'}{width=0.5
height=0.5}{qrcode}} % 4th QR Code
\\rput[Bl](0.7,19.26){'${texName}'}' >> latexTestScript.tex

num=$(($num+1))

fi
```

```
if [ $length -gt $num ]
then

texName=$(echo ${names[$num]}  | awk '{gsub( /[#$%&_{}]/, "\\\\&");
print $0}')

echo '\\rput(0.7,19.98){\psbarcode{'${names[$num]}'}{width=0.5
height=0.5}{qrcode}} % 4th QR Code
\\rput[Bl](0.7,21.38){'${texName}'}' >> latexTestScript.tex

num=$(($num+1))

fi

if [ $length -gt $num ]
then

texName=$(echo ${names[$num]}  | awk '{gsub( /[#$%&_{}]/, "\\\\&");
print $0}')

echo '\\rput(0.7,22.1){\psbarcode{'${names[$num]}'}{width=0.5
height=0.5}{qrcode}} % 4th QR Code
\\rput[Bl](0.7,23.5){'${texName}'}' >> latexTestScript.tex

num=$(($num+1))

fi

if [ $length -gt $num ]
then

texName=$(echo ${names[$num]}  | awk '{gsub( /[#$%&_{}]/, "\\\\&");
print $0}')

echo '\\rput(0.7,24.22){\psbarcode{'${names[$num]}'}{width=0.5
height=0.5}{qrcode}} % 4th QR Code
\\rput[Bl](0.7,25.62){'${texName}'}' >> latexTestScript.tex

num=$(($num+1))

fi

if [ $length -gt $num ]
then

texName=$(echo ${names[$num]}  | awk '{gsub( /[#$%&_{}]/, "\\\\&");
print $0}')

echo '\\rput(0.7,26.34){\psbarcode{'${names[$num]}'}{width=0.5
height=0.5}{qrcode}} % 4th QR Code
\\rput[Bl](0.7,27.74){'${texName}'}' >> latexTestScript.tex

num=$(($num+1))
```

```
fi

##column2
if [ $length -gt $num ]
then

texName=$(echo ${names[$num]}  | awk '{gsub( /[#$%&_{}]/, "\\\\&");
print $0}')

echo '\\rput(4.5,0.9){\psbarcode{'${names[$num]}'}{width=0.5
height=0.5}{qrcode}} % 4th QR Code
\\rput[Bl](4.5,2.3){'${texName}'}' >> latexTestScript.tex

num=$(($num+1))

fi

if [ $length -gt $num ]
then

texName=$(echo ${names[$num]}  | awk '{gsub( /[#$%&_{}]/, "\\\\&");
print $0}')

echo '\\rput(4.5,3.02){\psbarcode{'${names[$num]}'}{width=0.5
height=0.5}{qrcode}} % 4th QR Code
\\rput[Bl](4.5,4.42){'${texName}'}' >> latexTestScript.tex

num=$(($num+1))

fi

if [ $length -gt $num ]
then

texName=$(echo ${names[$num]}  | awk '{gsub( /[#$%&_{}]/, "\\\\&");
print $0}')

echo '\\rput(4.5,5.14){\psbarcode{'${names[$num]}'}{width=0.5
height=0.5}{qrcode}} % 4th QR Code
\\rput[Bl](4.5,6.54){'${texName}'}' >> latexTestScript.tex

num=$(($num+1))

fi

if [ $length -gt $num ]
then

texName=$(echo ${names[$num]}  | awk '{gsub( /[#$%&_{}]/, "\\\\&");
print $0}')

echo '\\rput(4.5,7.26){\psbarcode{'${names[$num]}'}{width=0.5
height=0.5}{qrcode}} % 4th QR Code
\\rput[Bl](4.5,8.66){'${texName}'}' >> latexTestScript.tex
```

```
num=$(($num+1))

fi

if [ $length -gt $num ]
then

texName=$(echo ${names[$num]}  | awk '{gsub( /[#$%&_{}]/, "\\\\&");
print $0}')

echo '\\rput(4.5,9.38){\psbarcode{'${names[$num]}'}{width=0.5
height=0.5}{qrcode}} % 4th QR Code
\\rput[Bl](4.5,10.78){'${texName}'}' >> latexTestScript.tex

num=$(($num+1))

fi

if [ $length -gt $num ]
then

texName=$(echo ${names[$num]}  | awk '{gsub( /[#$%&_{}]/, "\\\\&");
print $0}')

echo '\\rput(4.5,11.5){\psbarcode{'${names[$num]}'}{width=0.5
height=0.5}{qrcode}} % 4th QR Code
\\rput[Bl](4.5,12.9){'${texName}'}' >> latexTestScript.tex

num=$(($num+1))

fi


if [ $length -gt $num ]
then

texName=$(echo ${names[$num]}  | awk '{gsub( /[#$%&_{}]/, "\\\\&");
print $0}')

echo '\\rput(4.5,13.62){\psbarcode{'${names[$num]}'}{width=0.5
height=0.5}{qrcode}} % 4th QR Code
\\rput[Bl](4.5,15.02){'${texName}'}' >> latexTestScript.tex

num=$(($num+1))

fi

if [ $length -gt $num ]
then

texName=$(echo ${names[$num]}  | awk '{gsub( /[#$%&_{}]/, "\\\\&");
print $0}')
```

```
echo '\\rput(4.5,15.74){\psbarcode{'${names[$num]}'}{width=0.5
height=0.5}{qrcode}} % 4th QR Code
\\rput[Bl](4.5,17.14){'${texName}'}' >> latexTestScript.tex

num=$(($num+1))

fi


if [ $length -gt $num ]
then

texName=$(echo ${names[$num]}  | awk '{gsub( /[#$%&_{}]/, "\\\\&");
print $0}')

echo '\\rput(4.5,17.86){\psbarcode{'${names[$num]}'}{width=0.5
height=0.5}{qrcode}} % 4th QR Code
\\rput[Bl](4.5,19.26){'${texName}'}' >> latexTestScript.tex

num=$(($num+1))

fi

if [ $length -gt $num ]
then

texName=$(echo ${names[$num]}  | awk '{gsub( /[#$%&_{}]/, "\\\\&");
print $0}')

echo '\\rput(4.5,19.98){\psbarcode{'${names[$num]}'}{width=0.5
height=0.5}{qrcode}} % 4th QR Code
\\rput[Bl](4.5,21.38){'${texName}'}' >> latexTestScript.tex

num=$(($num+1))

fi

if [ $length -gt $num ]
then

texName=$(echo ${names[$num]}  | awk '{gsub( /[#$%&_{}]/, "\\\\&");
print $0}')

echo '\\rput(4.5,22.1){\psbarcode{'${names[$num]}'}{width=0.5
height=0.5}{qrcode}} % 4th QR Code
\\rput[Bl](4.5,23.5){'${texName}'}' >> latexTestScript.tex

num=$(($num+1))

fi

if [ $length -gt $num ]
then
```

```
texName=$(echo ${names[$num]}  | awk '{gsub( /[#$%&_{}]/, "\\\\&");
print $0}')

echo '\\rput(4.5,24.22){\psbarcode{'${names[$num]}'}{width=0.5
height=0.5}{qrcode}} % 4th QR Code
\\rput[Bl](4.5,25.62){'${texName}'}' >> latexTestScript.tex

num=$(($num+1))

fi

if [ $length -gt $num ]
then

texName=$(echo ${names[$num]}  | awk '{gsub( /[#$%&_{}]/, "\\\\&");
print $0}')

echo '\\rput(4.5,26.34){\psbarcode{'${names[$num]}'}{width=0.5
height=0.5}{qrcode}} % 4th QR Code
\\rput[Bl](4.5,27.74){'${texName}'}' >> latexTestScript.tex

num=$(($num+1))

fi

##column3


if [ $length -gt $num ]
then

texName=$(echo ${names[$num]}  | awk '{gsub( /[#$%&_{}]/, "\\\\&");
print $0}')

echo '\\rput(8.3,0.9){\psbarcode{'${names[$num]}'}{width=0.5
height=0.5}{qrcode}} % 4th QR Code
\\rput[Bl](8.3,2.3){'${texName}'}' >> latexTestScript.tex

num=$(($num+1))

fi


if [ $length -gt $num ]
then

texName=$(echo ${names[$num]}  | awk '{gsub( /[#$%&_{}]/, "\\\\&");
print $0}')

echo '\\rput(8.3,3.02){\psbarcode{'${names[$num]}'}{width=0.5
height=0.5}{qrcode}} % 4th QR Code
\\rput[Bl](8.3,4.42){'${texName}'}' >> latexTestScript.tex

num=$(($num+1))
```

```
fi

if [ $length -gt $num ]
then

texName=$(echo ${names[$num]}  | awk '{gsub( /[#$%&_{}]/, "\\\\&");
print $0}')

echo '\\rput(8.3,5.14){\psbarcode{'${names[$num]}'}{width=0.5
height=0.5}{qrcode}} % 4th QR Code
\\rput[Bl](8.3,6.54){'${texName}'}' >> latexTestScript.tex

num=$(($num+1))

fi


if [ $length -gt $num ]
then

texName=$(echo ${names[$num]}  | awk '{gsub( /[#$%&_{}]/, "\\\\&");
print $0}')

echo '\\rput(8.3,7.26){\psbarcode{'${names[$num]}'}{width=0.5
height=0.5}{qrcode}} % 4th QR Code
\\rput[Bl](8.3,8.66){'${texName}'}' >> latexTestScript.tex

num=$(($num+1))

fi

if [ $length -gt $num ]
then

texName=$(echo ${names[$num]}  | awk '{gsub( /[#$%&_{}]/, "\\\\&");
print $0}')

echo '\\rput(8.3,9.38){\psbarcode{'${names[$num]}'}{width=0.5
height=0.5}{qrcode}} % 4th QR Code
\\rput[Bl](8.3,10.78){'${texName}'}' >> latexTestScript.tex

num=$(($num+1))

fi

if [ $length -gt $num ]
then

texName=$(echo ${names[$num]}  | awk '{gsub( /[#$%&_{}]/, "\\\\&");
print $0}')

echo '\\rput(8.3,11.5){\psbarcode{'${names[$num]}'}{width=0.5
height=0.5}{qrcode}} % 4th QR Code
```

```
\\rput[Bl](8.3,12.9){'${texName}'}' >> latexTestScript.tex

num=$(($num+1))

fi

if [ $length -gt $num ]
then

texName=$(echo ${names[$num]}  | awk '{gsub( /[#$%&_{}]/, "\\\\&");
print $0}')

echo '\\rput(8.3,13.62){\psbarcode{'${names[$num]}'}{width=0.5
height=0.5}{qrcode}} % 4th QR Code
\\rput[Bl](8.3,15.02){'${texName}'}' >> latexTestScript.tex

num=$(($num+1))

fi


if [ $length -gt $num ]
then

texName=$(echo ${names[$num]}  | awk '{gsub( /[#$%&_{}]/, "\\\\&");
print $0}')

echo '\\rput(8.3,15.74){\psbarcode{'${names[$num]}'}{width=0.5
height=0.5}{qrcode}} % 4th QR Code
\\rput[Bl](8.3,17.14){'${texName}'}' >> latexTestScript.tex

num=$(($num+1))

fi

if [ $length -gt $num ]
then

texName=$(echo ${names[$num]}  | awk '{gsub( /[#$%&_{}]/, "\\\\&");
print $0}')

echo '\\rput(8.3,17.86){\psbarcode{'${names[$num]}'}{width=0.5
height=0.5}{qrcode}} % 4th QR Code
\\rput[Bl](8.3,19.26){'${texName}'}' >> latexTestScript.tex

num=$(($num+1))

fi

if [ $length -gt $num ]
then

texName=$(echo ${names[$num]}  | awk '{gsub( /[#$%&_{}]/, "\\\\&");
print $0}')
```

```
echo '\\rput(8.3,19.98){\psbarcode{'${names[$num]}'}{width=0.5
height=0.5}{qrcode}} % 4th QR Code
\\rput[Bl](8.3,21.38){'${texName}'}' >> latexTestScript.tex

num=$(($num+1))

fi

if [ $length -gt $num ]
then

texName=$(echo ${names[$num]}  | awk '{gsub( /[#$%&_{}]/, "\\\\&");
print $0}')

echo '\\rput(8.3,22.1){\psbarcode{'${names[$num]}'}{width=0.5
height=0.5}{qrcode}} % 4th QR Code
\\rput[Bl](8.3,23.5){'${texName}'}' >> latexTestScript.tex

num=$(($num+1))

fi

if [ $length -gt $num ]
then

texName=$(echo ${names[$num]}  | awk '{gsub( /[#$%&_{}]/, "\\\\&");
print $0}')

echo '\\rput(8.3,24.22){\psbarcode{'${names[$num]}'}{width=0.5
height=0.5}{qrcode}} % 4th QR Code
\\rput[Bl](8.3,25.62){'${texName}'}' >> latexTestScript.tex

num=$(($num+1))

fi

if [ $length -gt $num ]
then

texName=$(echo ${names[$num]}  | awk '{gsub( /[#$%&_{}]/, "\\\\&");
print $0}')

echo '\\rput(8.3,26.34){\psbarcode{'${names[$num]}'}{width=0.5
height=0.5}{qrcode}} % 4th QR Code
\\rput[Bl](8.3,27.74){'${texName}'}' >> latexTestScript.tex

num=$(($num+1))

fi

## clomn4
if [ $length -gt $num ]
then
```

```
texName=$(echo ${names[$num]}  | awk '{gsub( /[#$%&_{}]/, "\\\\&");
print $0}')

echo '\\rput(12.1,0.9){\psbarcode{'${names[$num]}'}{width=0.5
height=0.5}{qrcode}} % 4th QR Code
\\rput[Bl](12.1,2.3){'${texName}'}' >> latexTestScript.tex

num=$(($num+1))

fi

if [ $length -gt $num ]
then

texName=$(echo ${names[$num]}  | awk '{gsub( /[#$%&_{}]/, "\\\\&");
print $0}')

echo '\\rput(12.1,3.02){\psbarcode{'${names[$num]}'}{width=0.5
height=0.5}{qrcode}} % 4th QR Code
\\rput[Bl](12.1,4.42){'${texName}'}' >> latexTestScript.tex

num=$(($num+1))

fi

if [ $length -gt $num ]
then

texName=$(echo ${names[$num]}  | awk '{gsub( /[#$%&_{}]/, "\\\\&");
print $0}')

echo '\\rput(12.1,5.14){\psbarcode{'${names[$num]}'}{width=0.5
height=0.5}{qrcode}} % 4th QR Code
\\rput[Bl](12.1,6.54){'${texName}'}' >> latexTestScript.tex

num=$(($num+1))

fi


if [ $length -gt $num ]
then

texName=$(echo ${names[$num]}  | awk '{gsub( /[#$%&_{}]/, "\\\\&");
print $0}')

echo '\\rput(12.1,7.26){\psbarcode{'${names[$num]}'}{width=0.5
height=0.5}{qrcode}} % 4th QR Code
\\rput[Bl](12.1,8.66){'${texName}'}' >> latexTestScript.tex

num=$(($num+1))

fi
```

```
if [ $length -gt $num ]
then

texName=$(echo ${names[$num]}  | awk '{gsub( /[#$%&_{}]/, "\\\\&");
print $0}')

echo '\\rput(12.1,9.38){\psbarcode{'${names[$num]}'}{width=0.5
height=0.5}{qrcode}} % 4th QR Code
\\rput[Bl](12.1,10.78){'${texName}'}' >> latexTestScript.tex

num=$(($num+1))

fi

if [ $length -gt $num ]
then

texName=$(echo ${names[$num]}  | awk '{gsub( /[#$%&_{}]/, "\\\\&");
print $0}')

echo '\\rput(12.1,11.5){\psbarcode{'${names[$num]}'}{width=0.5
height=0.5}{qrcode}} % 4th QR Code
\\rput[Bl](12.1,12.9){'${texName}'}' >> latexTestScript.tex

num=$(($num+1))

fi

if [ $length -gt $num ]
then

texName=$(echo ${names[$num]}  | awk '{gsub( /[#$%&_{}]/, "\\\\&");
print $0}')

echo '\\rput(12.1,13.62){\psbarcode{'${names[$num]}'}{width=0.5
height=0.5}{qrcode}} % 4th QR Code
\\rput[Bl](12.1,15.02){'${texName}'}' >> latexTestScript.tex

num=$(($num+1))

fi


if [ $length -gt $num ]
then

texName=$(echo ${names[$num]}  | awk '{gsub( /[#$%&_{}]/, "\\\\&");
print $0}')

echo '\\rput(12.1,15.74){\psbarcode{'${names[$num]}'}{width=0.5
height=0.5}{qrcode}} % 4th QR Code
\\rput[Bl](12.1,17.14){'${texName}'}' >> latexTestScript.tex
```

```
num=$(($num+1))

fi

if [ $length -gt $num ]
then

texName=$(echo ${names[$num]}  | awk '{gsub( /[#$%&_{}]/, "\\\\&");
print $0}')

echo '\\rput(12.1,17.86){\psbarcode{'${names[$num]}'}{width=0.5
height=0.5}{qrcode}} % 4th QR Code
\\rput[Bl](12.1,19.26){'${texName}'}' >> latexTestScript.tex

num=$(($num+1))

fi

if [ $length -gt $num ]
then

texName=$(echo ${names[$num]}  | awk '{gsub( /[#$%&_{}]/, "\\\\&");
print $0}')

echo '\\rput(12.1,19.98){\psbarcode{'${names[$num]}'}{width=0.5
height=0.5}{qrcode}} % 4th QR Code
\\rput[Bl](12.1,21.38){'${texName}'}' >> latexTestScript.tex

num=$(($num+1))

fi


if [ $length -gt $num ]
then

texName=$(echo ${names[$num]}  | awk '{gsub( /[#$%&_{}]/, "\\\\&");
print $0}')

echo '\\rput(12.1,22.1){\psbarcode{'${names[$num]}'}{width=0.5
height=0.5}{qrcode}} % 4th QR Code
\\rput[Bl](12.1,23.5){'${texName}'}' >> latexTestScript.tex

num=$(($num+1))

fi

if [ $length -gt $num ]
then

texName=$(echo ${names[$num]}  | awk '{gsub( /[#$%&_{}]/, "\\\\&");
print $0}')

echo '\\rput(12.1,24.22){\psbarcode{'${names[$num]}'}{width=0.5
```

```
height=0.5}{qrcode}} % 4th QR Code
\\rput[Bl](12.1,25.62){'${texName}'}' >> latexTestScript.tex

num=$(($num+1))

fi

if [ $length -gt $num ]
then

texName=$(echo ${names[$num]}  | awk '{gsub( /[#$%&_{}]/, "\\\\&");
print $0}')

echo '\\rput(12.1,26.34){\psbarcode{'${names[$num]}'}{width=0.5
height=0.5}{qrcode}} % 4th QR Code
\\rput[Bl](12.1,27.74){'${texName}'}' >> latexTestScript.tex

num=$(($num+1))

fi


##column5
if [ $length -gt $num ]
then

texName=$(echo ${names[$num]}  | awk '{gsub( /[#$%&_{}]/, "\\\\&");
print $0}')

echo '\\rput(15.9,0.9){\psbarcode{'${names[$num]}'}{width=0.5
height=0.5}{qrcode}} % 4th QR Code
\\rput[Bl](15.9,2.3){'${texName}'}' >> latexTestScript.tex

num=$(($num+1))

fi

if [ $length -gt $num ]
then

texName=$(echo ${names[$num]}  | awk '{gsub( /[#$%&_{}]/, "\\\\&");
print $0}')

echo '\\rput(15.9,3.02){\psbarcode{'${names[$num]}'}{width=0.5
height=0.5}{qrcode}} % 4th QR Code
\\rput[Bl](15.9,4.42){'${texName}'}' >> latexTestScript.tex

num=$(($num+1))

fi

if [ $length -gt $num ]
then
```

```
texName=$(echo ${names[$num]}  | awk '{gsub( /[#$%&_{}]/, "\\\\&");
print $0}')

echo '\\rput(15.9,5.14){\psbarcode{'${names[$num]}'}{width=0.5
height=0.5}{qrcode}} % 4th QR Code
\\rput[Bl](15.9,6.54){'${texName}'}' >> latexTestScript.tex

num=$(($num+1))

fi


if [ $length -gt $num ]
then

texName=$(echo ${names[$num]}  | awk '{gsub( /[#$%&_{}]/, "\\\\&");
print $0}')

echo '\\rput(15.9,7.26){\psbarcode{'${names[$num]}'}{width=0.5
height=0.5}{qrcode}} % 4th QR Code
\\rput[Bl](15.9,8.66){'${texName}'}' >> latexTestScript.tex

num=$(($num+1))

fi

if [ $length -gt $num ]
then

texName=$(echo ${names[$num]}  | awk '{gsub( /[#$%&_{}]/, "\\\\&");
print $0}')

echo '\\rput(15.9,9.38){\psbarcode{'${names[$num]}'}{width=0.5
height=0.5}{qrcode}} % 4th QR Code
\\rput[Bl](15.9,10.78){'${texName}'}' >> latexTestScript.tex

num=$(($num+1))

fi

if [ $length -gt $num ]
then

texName=$(echo ${names[$num]}  | awk '{gsub( /[#$%&_{}]/, "\\\\&");
print $0}')

echo '\\rput(15.9,11.5){\psbarcode{'${names[$num]}'}{width=0.5
height=0.5}{qrcode}} % 4th QR Code
\\rput[Bl](15.9,12.9){'${texName}'}' >> latexTestScript.tex

num=$(($num+1))

fi
```

```
if [ $length -gt $num ]
then

texName=$(echo ${names[$num]}  | awk '{gsub( /[#$%&_{}]/, "\\\\&");
print $0}')

echo '\\rput(15.9,13.62){\psbarcode{'${names[$num]}'}{width=0.5
height=0.5}{qrcode}} % 4th QR Code
\\rput[Bl](15.9,15.02){'${texName}'}' >> latexTestScript.tex

num=$(($num+1))

fi


if [ $length -gt $num ]
then

texName=$(echo ${names[$num]}  | awk '{gsub( /[#$%&_{}]/, "\\\\&");
print $0}')

echo '\\rput(15.9,15.74){\psbarcode{'${names[$num]}'}{width=0.5
height=0.5}{qrcode}} % 4th QR Code
\\rput[Bl](15.9,17.14){'${texName}'}' >> latexTestScript.tex

num=$(($num+1))

fi

if [ $length -gt $num ]
then

texName=$(echo ${names[$num]}  | awk '{gsub( /[#$%&_{}]/, "\\\\&");
print $0}')

echo '\\rput(15.9,17.86){\psbarcode{'${names[$num]}'}{width=0.5
height=0.5}{qrcode}} % 4th QR Code
\\rput[Bl](15.9,19.26){'${texName}'}' >> latexTestScript.tex

num=$(($num+1))

fi

if [ $length -gt $num ]
then

texName=$(echo ${names[$num]}  | awk '{gsub( /[#$%&_{}]/, "\\\\&");
print $0}')

echo '\\rput(15.9,19.98){\psbarcode{'${names[$num]}'}{width=0.5
height=0.5}{qrcode}} % 4th QR Code
\\rput[Bl](15.9,21.38){'${texName}'}' >> latexTestScript.tex

num=$(($num+1))
```

```
fi


if [ $length -gt $num ]
then

texName=$(echo ${names[$num]}  | awk '{gsub( /[#$%&_{}]/, "\\\\&");
print $0}')

echo '\\rput(15.9,22.1){\psbarcode{'${names[$num]}'}{width=0.5
height=0.5}{qrcode}} % 4th QR Code
\\rput[Bl](15.9,23.5){'${texName}'}' >> latexTestScript.tex

num=$(($num+1))

fi

if [ $length -gt $num ]
then

texName=$(echo ${names[$num]}  | awk '{gsub( /[#$%&_{}]/, "\\\\&");
print $0}')

echo '\\rput(15.9,24.22){\psbarcode{'${names[$num]}'}{width=0.5
height=0.5}{qrcode}} % 4th QR Code
\\rput[Bl](15.9,25.62){'${texName}'}' >> latexTestScript.tex

num=$(($num+1))

fi

if [ $length -gt $num ]
then

texName=$(echo ${names[$num]}  | awk '{gsub( /[#$%&_{}]/, "\\\\&");
print $0}')

echo '\\rput(15.9,26.34){\psbarcode{'${names[$num]}'}{width=0.5
height=0.5}{qrcode}} % 4th QR Code
\\rput[Bl](15.9,27.74){'${texName}'}' >> latexTestScript.tex

num=$(($num+1))

fi

## End of page

echo "\end{pspicture}

\end{center}" >> latexTestScript.tex

done
```

```
## End of document

echo "\end{document}" >> latexTestScript.tex

## Submit script

mv latexTestScript.tex $output #Re-name .tex script to output name

pdflatex -shell-escape $output # Run .tex script
```
