## Supplemental_File_2 for "Cultivar-specific fungicide resistance emerges during a growing season in field populations of *Zymoseptoria tritici*"

```
######Talas et al.
2024##############################################################
###Cultivar-specific fungicide resistance emerges during a growing
season in field populations ####of Zymoseptoria tritici###
////////////////////////////////////////////////////////////////////
////////////////////////////////////////////////////////////////////
////////////////////////////////////

dir = getDirectory("Choose a Directory ");

list = getFileList(dir);
for (ph=0; ph<list.length; ph++) {
image_path=dir + list[ph];

open(image_path);


run("Barcode Codec");

temp_path=dir+"temp.txt";

saveAs("Text", temp_path);
    close();
    close("Decoded Text");

code=File.openAsString(temp_path);
File.delete(temp_path);


data=split(code, "");
image_name=data[0];
dir_name=split(image_name,"_");
dir_name=dir_name[0];


new_folder=dir+dir_name;

File.makeDirectory(new_folder);
File.makeDirectory(new_folder+"/Photos");
File.makeDirectory(new_folder+"/Labels");
File.makeDirectory(new_folder+"/Masks");
File.makeDirectory(new_folder+"/Results");





File.rename(image_path,new_folder+"/"+"Photos"+"/"+image_name+".jpg"
);

               }
```

```
run("Close");
run("Close All");
```
