## Supplemental_File_3 for "Cultivar-specific fungicide resistance emerges during a growing season in field populations of *Zymoseptoria tritici*"

```
////////////////////////////////////////////////////////////////////
////////////////////////////////////////////////////////////////////
/////////////////////////////////////////
### this macro was developed based on 1. https://doi.org/10.1002/
bmb.21139. ####2. Lendenmann et al. 2015############################
////////////////////////////////////////////////////////////////////
////////////////////////////////////////////////////////////////////
///////////////////////////////////////
######Talas et al. 2024. Cultivar-specific fungicide resistance
emerges during a growing season ###in field populations of
Zymoseptoria
##tritici###########################################################
###
////////////////////////////////////////////////////////////////////
////////////////////////////////////////////////////////////////////
////////////////////////////////////


      dir = getDirectory("Choose a Directory ");

      list = getFileList(dir+"Photos");
      print("list");

      for (ph=0; ph<list.length; ph++) {

                         open(dir + "Photos/"+ list[ph]);



                          makeRectangle(831, 1389, 1017, 1506);
                          run("Crop");
                          run("Flip Horizontally");
                          run("Rotate 90 Degrees Left");

                           if (bitDepth != 24) {
                            exit("This macro requires an RGB
image");
                             }

                          nc=12;
                          nr=8;
                          xo=164.5;
                          yo=140;
                          xf=1334;
                          yf=878;
                          csize=72.50;


                             function create_grid_dialog() {
                             Dialog.create("Grid Parameters");
                             Dialog.addNumber("Number of Columns
(max 12):", nc);
                             Dialog.addNumber("Number of Rows (max
8):", nr);
```

```
                             Dialog.addNumber("Center of well A1: X
origin (in pixels, left equals zero):", xo);
                             Dialog.addNumber("Center of well A1: Y
origin (in pixels, up equals zero):", yo);
                             Dialog.addNumber("Center of well H12: X
end (in pixels):", xf);
                             Dialog.addNumber("Center of well H12: Y
end (in pixels):", yf);
                             Dialog.addNumber("Circle Size (Diameter
in pixels):", csize);
                             Dialog.show();
                             }

                            function validate_grid_parameters(nc,
nr, xo, yo, xf, yf, csize) {
                            if (nc < 1 || nc > 12) {
                            exit("The number of columns must be
between 1 and 12");
                             }
                             if (nr < 1 || nr > 8) {
                            exit("The number of rows must be between
1 and 8");
                             }
                            if (xo < 0) {
                            exit("The X origin cannot be negative");
                             }
                             if (yo < 0) {
                             exit ("The Y origin cannot be
negative");
                             }
                            if (xf < xo) {
                            exit("The X end cannot be lower than the
X origin");
                            }
                            if (yf < yo) {
                            exit("The Y end cannot be lower than the
Y origin");
                            }
                           if (csize < 1) {
                           exit("The Diameter should be at least one
pixel");
                           }
                         }

                           function mark_selections(nr, nc, x, y,
csepx, csepy, csize) {
                           for (i = 0; i < nr; i++) {
                           for (j = 0; j < nc; j++) {
                           makeOval(x+j*csepx, y+i*csepy, csize,
csize);
                           run("Add Selection...", "stroke=yellow
width=1");
```

```
                }
                }
            }

                csepx = (xf - xo) / 11;
                csepy = (yf - yo) / 7;


                x = xo - (csize / 2);
                y = yo - (csize / 2);
                csmallsize = csize / sqrt(3);

                mark_selections(nr, nc, x, y, csepx,
csepy, csize);
                run("Labels...", "color=black font=24
show labels");


                 index = 0;
                 for (i = 0; i < nr; i++) {
                 row = substring("ABCDEFGH", i, i+1);
                 for (j = 0; j < nc; j++) {
                 Overlay.getBounds(index, x_well, y_well,
w, h);
                 index++;

                 makeOval(x+(j-0.5)*csepx, y+
(i-0.5)*csepy, csmallsize, csmallsize);
                 getStatistics(area, mean);
                 b1 = mean;

                 makeOval(x+(j-0.5)*csepx, y+
(i+0.5)*csepy, csmallsize, csmallsize);
                 getStatistics(area, mean);
                 b2 = mean;

                 makeOval(x+(j+0.5)*csepx, y+
(i-0.5)*csepy, csmallsize, csmallsize);
                 getStatistics(area, mean);
                 b3 = mean;

                  makeOval(x+(j+0.5)*csepx, y+
(i+0.5)*csepy, csmallsize, csmallsize);
                  getStatistics(area, mean);
                  b4 = mean;

                   b12min = minOf(b1,b2);
                   b34min = minOf(b3,b4);
                   b14min = minOf(b12min,b34min);
                   bav = ((b1+b2+b3+b4) - b14min) / 3;
```

```
                                makeOval(x_well, y_well, w, h);
                                getStatistics(area, mean);
                                intensity = mean;

                                run("Measure");
                                setResult("Row", nResults-1, row);
                                setResult("Column", nResults-1, j+1);

                                 abuncorr = -(log(intensity/255)/
log(10));
                                 setResult("Auncorr", nResults-1,
abuncorr);

                                 ablank = -(log(bav/255)/log(10));
                                 setResult("Ablank", nResults-1,
ablank);

                                 abcorr = abuncorr - ablank;
                                 setResult("Acorr", nResults-1,
abcorr);

                                 updateResults();


                                 }
                           }


                                 saveAs("Jpeg", dir+ "Labels/"+
list[ph]);
                                 run("Remove Overlay");
                                 saveAs("Jpeg", dir+
"Masks/"+list[ph]);

                                 selectWindow("Results");
                                 name=replace(list[ph],".jpg",".txt");
                                 saveAs("Text", dir+ "Results/" +
name);
                                 close();
                                 selectWindow("Results");

       }

       run("Close");
```
